# The role of petal transpiration in floral humidity generation

**DOI:** 10.1101/2021.04.06.438589

**Authors:** Michael J. M. Harrap, Sean A. Rands

## Abstract

Floral humidity, an area of elevated humidity in the headspace of flowers, has been detected across angiosperms and may function as a pollinator cue for insect pollinators. It is believed floral humidity is produced predominantly through a combination of evaporation of both liquid nectar and transpirational water loss from the flower. However, the role of transpiration in floral humidity generation has not been tested and is largely inferred by continued humidity production when nectar is removed from flowers. Understanding the extent that transpiration contributes to floral humidity has important implications for understanding the function of floral humidity. We test whether transpiration contributes to the floral humidity generation of two species previously identified to produce elevated floral humidity, *Calystegia silvatica* and *Eschscholzia californica*. Floral humidity production of flowers that underwent an antitranspirant treatment, petrolatum gel which blocks transpiration from treated tissues, is compared to flowers that did not receive such treatments. Gel treatments reduced floral humidity production to approximately a third of that produced by untreated flowers in *C. silvatica*, and half of that in *E. californica*. This confirms, the previously untested, inferences that transpiration has a large contribution to floral humidity generation and that this contribution may vary between species.

**HIGHLIGHT:** We confirm, the previously untested, inferences that transpiration has a large contribution to floral humidity generation and show that this contribution may vary between species.

## INTRODUCTION

Floral humidity, an area of elevated humidity in the headspace of the flower (relative to the environment) has been detected in several flower species across different families (Harrap et al., 2020). Flower species have been found to vary in the intensity of floral humidity produced (the difference in humidity between the floral headspace and the environment) and in the humidity structure (the shape and location of elevated humidity within the flower headspace). Floral humidity may have important influences on flower function and fitness. Pollinators use different floral signalling modalities to inform foraging decisions (Leonard et al., 2011, 2012; Leonard & Masek, 2014; Raguso, 2004) and floral humidity, as part of a multimodal floral display, may influence these. Hawkmoths (von Arx et al., 2012), bumblebees (Harrap et al., 2021) and flies (Nordström et al., 2017) have innate preferences for flowers that produce higher floral humidity intensities when they are able to choose between flowers producing differing intensities. Furthermore, floral humidity differences between flowers, regardless of whether elevated rewards are associated with higher floral humidity production or not, can aid bumblebee learning of rewarding flowers (Harrap et al., 2021). By influencing floral preferences and learning, floral humidity may affect foraging success of naïve and experienced pollinators (Raine & Chittka, 2008) and will in turn influence visitation rates of pollinators and thus pollen receipt and export (Ashman et al., 2004; Schiestl & Johnson, 2013). Humidity production by flowers, perhaps alongside production from vegetative tissue, may also affect patch-level foraging decisions by pollinators, allowing them to locate areas of elevated foliage (where suitable forage is more likely to be) resulting in similar impacts on pollinator and plant fitness (Wolfin et al., 2018). Floral humidity may have further influences on plant fitness that have not yet been directly investigated. Humidity conditions influence pollen water content, which in turn influences the viability and germination ability of pollen (Hase et al., 2006; Nepi et al., 2001). Floral humidity may therefore have an influence on pollen viability, perhaps maintaining an environment more suitable for pollen.

Floral humidity is believed to be generated through a combination of evaporation of liquid nectar of flowers and release of water vapour from the flower *via* transpiration, particularly from flower petals. However, other floral characteristics, such as floral structure, influence how humidity produced accumulates in the flower headspace (Harrap et al., 2020; von Arx et al., 2012). How flowers generate humidity has received little investigation, and this work has focused on the role of nectar evaporation. Removal of nectar from flowers of evening primrose *Oenothera caespitosa* resulted in a decrease in floral humidity production, and blocking nectaries also yielded similar results (von Arx et al., 2012), confirming the role of nectar evaporation in floral humidity generation. This is also supported by evidence that, within flower headspaces, proximity to nectaries is associated with increased humidity (Corbet, Unwin, et al., 1979; Corbet, Willmer, et al., 1979; Harrap et al., 2020; von Arx et al., 2012). Evidence of the contribution of transpiration to floral humidity is comparatively limited, and largely inferred. That nectar removal did not completely reduce floral humidity production in *O. caespitosa* indicates that there is an additional humidity source contributing to floral humidity (Harrap et al., 2020), and the most likely candidate for this is floral transpiration. Similarly, humidity detected from *O. caespitosa* was reduced when petal surfaces, but not nectar or nectaries, were shielded from sensors (von Arx et al., 2012), indicating petals contribute to floral humidity, likely *via* transpiration. However, there are other potential sources of floral humidity generation, such as the moist surfaces of reproductive structures. These may explain floral humidity production independent of nectar evaporation. Although the inference of transpiration’s role in floral humidity generation is logical, the contribution of transpiration to floral humidity generation has not been tested.

Petal permeability, and therefore floral transpiration rates, can be influenced by cuticle thickness, surface area and chemical composition (Buschhaus et al., 2015; Cheng et al., 2019; Guo et al., 2017; Hajibagheri et al., 1983). The presence, density and activity of petal stomata will also contribute to rates of floral transpiration (Hew et al., 1980; Huang et al., 2018; van Doorn, 1997; von Arx et al., 2012). Confirmation of the role of transpiration in floral humidity generation will confirm whether differences in such traits may help explain the diversity of floral humidity seen in angiosperms (Harrap et al., 2020). Environmental conditions (Gates, 1968; Jolliet & Bailey, 1992; Rawson et al., 1977; Schreiber, 2001) and plant daily cycles can affect transpiration (Simon et al., 2020), including floral transpiration cycles (Azad et al., 2007; Huang et al., 2018; Lü et al., 2011), so the extent to which floral humidity depends on transpiration may indicate potential for parallel changes in floral humidity with time and conditions, perhaps leading to changes in humidity cues and plant disease susceptibility. Furthermore, as floral humidity produced by transpiration will not be affected by nectar removal (von Arx et al., 2012), the role of transpiration in different species (particularly relative to nectar evaporation) may influence how well floral humidity cues indicate the temporary rewardlessness of the flower due to recent visits, influencing the extent that floral humidity can function as a directly ‘honest signal’ for pollinators (von Arx, 2013). Understanding the influence that floral transpiration has on floral humidity is therefore important for understanding both the function and evolution of floral humidity.

In this study, we demonstrate the contribution of petal transpiration to floral humidity generation in two flower species previously identified to produce elevated floral humidity. A simple antitranspirant treatment, petrolatum gel, is applied to the petals of flowers, serving to block transpirational water loss from petals. Humidity in the flower headspace is measured using robotic sampling techniques and compared between treated flowers and those of the same species which have not undergone this treatment. In this way, we evaluate floral humidity production with (untreated flowers) or without (gel-treated flowers) the contribution of petal transpiration.

## MATERIALS AND METHODS

### Collection and preparation of samples

The role of transpiration in floral humidity generation was tested in the flowers of giant bindweed *Calystegia silvatica* (Kit.) Griseb. and California poppy *Eschscholzia californica* Cham.. These species are appropriate choices for demonstrating the role of transpiration, as both produce larger amounts of floral humidity compared to other flower species (Harrap et al., 2020) comparable to levels which have been demonstrated to be able to influence pollinator foraging decisions (von Arx et al., 2012; von Arx, 2013; Wolfin et al., 2018; Harrap et al., 2021). *E. californica* has been identified (along with other poppies) to produce little in the way of nectar rewards (Hicks et al., 2016), and so nectar evaporation is unlikely to explain *E. californica* humidity production. *C. silvatica* produces more substantial nectar rewards (Baude et al., 2016), but is a useful study species as the Convolvulaceae are known to conduct large amounts of petal transpiration (Patiño & Grace, 2002).

Humidity sampling and treatments were also carried out on leaves of common ivy *Hedera helix* L.. Ivy leaves lack extrafloral nectaries or similar secretions (Vezza et al., 2006), so humidity produced by the leaf should be solely from transpiration sources *via* leaf stomata or epidermal tissue (von Arx et al., 2012). Consequently, leaf samples serve as a positive control for our gel treatment, confirming the extent that gel treatments block plant transpiration.

All plant material was collected from sites within walking distance of the University of Bristol main campus (51.45N 2.60W). Flowers were cut on the stem so no leaf remained on the cutting, but sepals were retained, and individual ivy leaves were cut with the majority of the petiole remaining attached. Immediately after cutting the stems attached to samples were stuck through a hole in the cap of a 24cm^3^ plastic horticulture tube, prefilled with water to within 2cm of the lid. Samples were transported upright to in a closed cardboard box. Flowers and leaves were only collected in dry conditions, so that no standing water from condensation or rain was present on flowers to influence the floral humidity measurements. Flowers and leaves were only used if they were fully open and did not show signs of age, disease, or damage.

Samples were collected to allow two samples (two flowers or two leaves) to be presented to the robot in pairs (spares were also collected to accommodate any breakage during treatment application). When selecting flowers and leaves that would be presented for sampling as ‘Gel’ and ‘Untreated’ sampling pairs (see below for treatment details), care was taken to collect flowers of approximately the same size, measured using a ruler across either the horizontal span of a flower, or the width of the leaf’s widest point. To control for size effects, flowers and leaves presented as sampling pairs of one ‘Gel’ and one ‘Untreated’ sample were within 10mm of their partner’s span. For sampled pairs of ‘Unhandled’ treatments, size was allowed to vary to capture as wide a range of the diversity as possible. Flowers and leaves presented for sampling in the same pair were always collected from the same site, so that flowers had grown in comparable conditions prior to being collected.

Prior to measurement, flowers were fixed in position, orientated vertically upwards, to prevent the measuring points of humidity transect differing with respect to the flower’s location due to the flower moving. Under natural condition, both species have vertically orientated flowers, although *C. silvatica* can have differing flower orientations dependent on where it grows. Thus, fixing the orientation of the flowers is unlikely to negatively affect the flowers. Flowers were fixed in position with a cylinder of rigid paper-card (Professional laser printer paper, Hewlett-Packard, Startbaan, Netherlands) fixed with tape (Scotch, St. Pauls, USA) to the outside of the horticultural tube, so the flowers rested flat facing vertically upwards (figure 1A and B). The size of the cylinder required varied with the individual flower and species being sampled, but normally protruded only a few centimetres above the tube’s lid and was of a comparable width to the tube itself. Care was taken not to affix this cylinder so that it constrained or compressed the flower.

**Figure 1:**
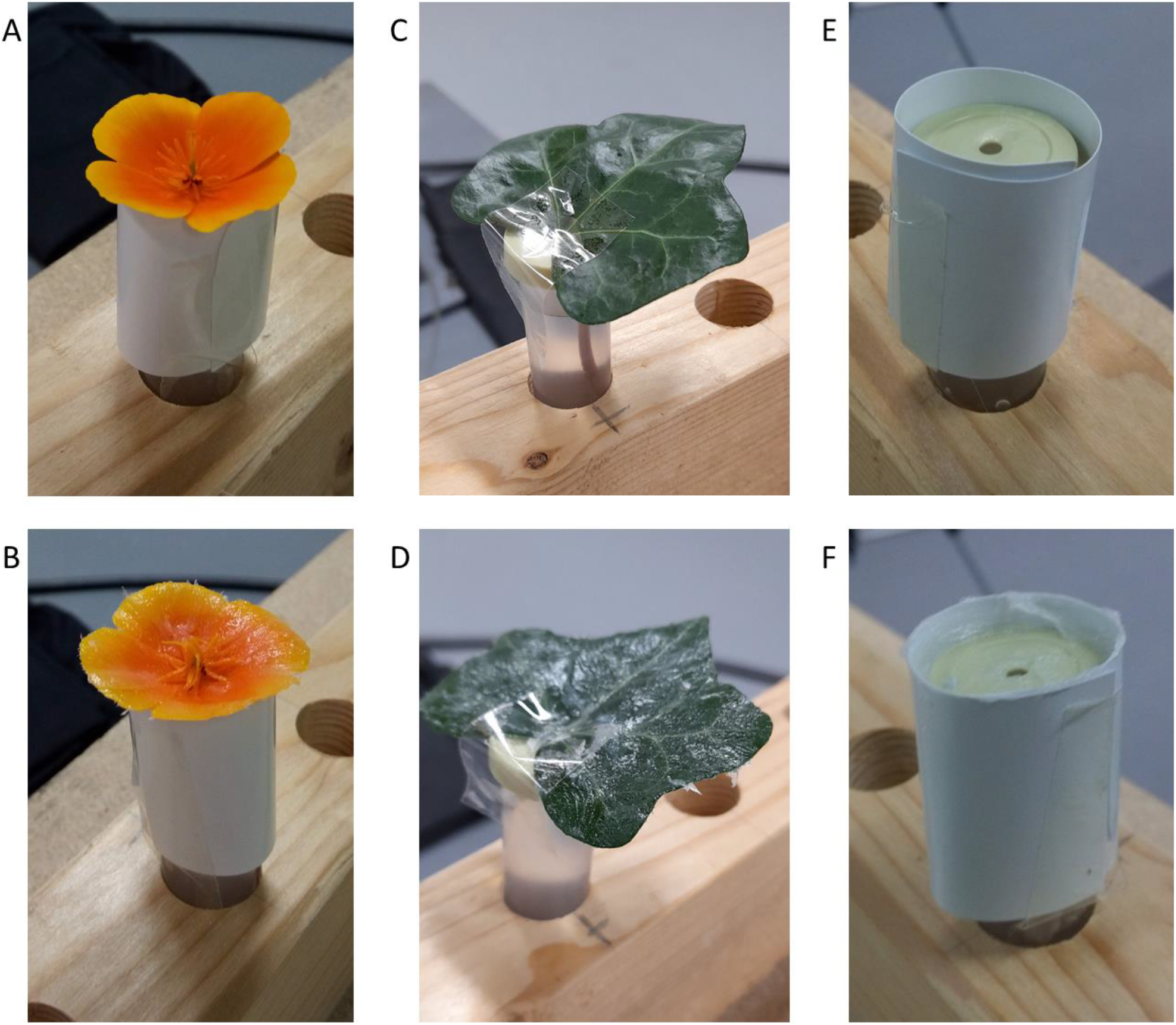
Prepared samples of: A, B, *Eschscholzia californica* flowers; C, D, *Hedera helix* leaves; E, F, tube control samples. A, C, E: ‘Untreated’ samples, where samples were handled as if gel were applied. B, D, F: ‘Gel’ treatment where gel was applied to sample surfaces. These samples are within a tube rack on a table before the robot, as they would be presented for humidity sampling.

Leaf samples could not be supported and orientated by card cylinders, and instead were secured to the horticultural tube with a section of tape after treatment application (described below). This tape was placed over the base of the leaf and the hole in the horticultural tube’s lid and continued down the side of the tube (figure 1C and D). This secured the leaves, which were angled slightly upwards relative to the horticultural tube cap. The irregular curving and shape of leaves meant that they did not always present a level surface, but this was accounted for when setting the ‘transect central point’ (see below).

### Preparation of control samples

Two further ‘control samples’ were conducted. These were the ‘Dry tube’ control, where a card support cylinder was attached around an empty horticultural tube (protruding 1cm above the lid), and the ‘Full tube’ control which was the same but with the tube filled with water to within 2cm of the lid, similar to leaf and flower samples (figure 1E and F). Controls served two purposes. First, when untreated, they allowed us to assess the extent that humidity differences detected between the focal and background probe are due to sources extraneous to the leaves or flowers placed within the tube. Evaporation of water *via* the hole in the lid could contribute to humidity difference (which we could measure with the ‘Full tube’ control). Similarly, random mixing of the air in the room can lead to uneven humidity environments and differences in humidity between the probes due to them differing in position (which could be measured with the ‘Dry tube’ control). Second, it is possible impurities in the petrolatum applied to samples evaporates or absorbs moisture allowing gel to affect humidity independently of blocking transpiration. By comparing the effect of gel treatments on the controls, we could evaluate whether the gel itself influences humidity.

### Antitranspirant treatments

Three treatments were conducted across flower, leaf and control samples. Flower samples underwent either the ‘Gel’, ‘Untreated’ or ‘Unhandled’ treatments (table 1). Leaf and control samples underwent only the ‘Gel’ and ‘Untreated’ treatments.

**Table 1:** A summary of the treatments applied to samples and the numbers of individuals of each sample type subjected to each treatment. Where ‘-’ is given samples were not subjected to that treatment. Bracketed values indicate the number of samples within that treatment that were also monitored for temperature differences.

| Treatment | Description | *Calystegia silvatica* | *Eschscholzia californica* | *Hedera helix* | Dry Tube Control | Full Tube Control |
| --- | --- | --- | --- | --- | --- | --- |
| Gel | Petrolatum gel is applied by hand over the abaxial and adaxial surface of flower petals and sepals and leaves, or the top of the tube lid in controls. | 25(7) | 25(6) | 8 | 15(2) | 19 |
| Untreated | Petrolatum gel is not applied. Samples are handled with dry hands as if gel were applied. | 25(7) | 25(6) | 8 | 15(2) | 19 |
| Unhandled | Petrolatum gel is not applied. Samples (flowers only) are not handled beyond collection and fixing of position in preparation for robot sampling. | 20(6) | 20(6) | - | - | - |

Samples receiving the ‘Gel’ treatment had a simple antitranspirant applied to their surface. In flowers receiving the ‘Gel’ treatment, petrolatum gel (‘Boots baby petroleum jelly’, Boots, Nottingham, UK) was applied evenly by hand over the abaxial and adaxial surface of petals, and over the flower sepals (figure 1B). Gel was not applied to floral reproductive structures. About the base of the petals’ upward (adaxial) surface where nectaries and reproductive structures were present, gel was applied as close as possible to these structures without covering them, meaning small areas of the petal near these structures remained untreated. This was to avoid the chance gel treatment may remove unconsidered contributions to floral humidity aside from petal (and sepal) transpiration such as nectar evaporation or release of moisture from reproductive structures. Similarly, in leaf samples that underwent the ‘Gel’ treatment, adaxial and abaxial surfaces of the leaf were covered with petrolatum gel (figure 1C), and in tube control samples gel was applied over the horticultural tube lid and around the protruding section of the inside of the card cylinder (figure 1F). Samples that underwent the ‘Untreated’ treatment had no antitranspirant gel applied on them, but the samples were handled as if it were being applied. Samples were handled with dry hands as described above, with similar spreading motions across the relevant surfaces associate with application of the gel. Only flower samples underwent the ‘Unhandled’ treatment, where no further handling of flowers was conducted following picking and fitting of card cylinder supports. Flowers in the ‘Unhandled’ treatment, being otherwise unmanipulated, allowed further evaluation of ‘normal’ floral humidity production. Comparing humidity production between the ‘Untreated’ and ‘Unhandled’ treatments allows us to account for the influences of the handling procedures involved with application of the gel antitransiprant when assessing the humidity production of treated flowers. Comparing ‘Gel’ treated to ‘Untreated’ flowers allows assessment of antitranspirant treatments on humidity production, accounting for the effects of handling. If at any time a sample broke during treatment application, it was replaced.

The humidity sampling sequence was conducted on a pair of samples of the same type (*i*.*e*. a pair of leaves or similar flowers, Dry tube controls, or Full tube controls). The pair of samples presented to the robot were prepared so that they were either: a sample that had undergone the ‘Gel’ treatment and sample that had undergone the ‘Untreated’ treatment; or two flower samples that had undergone the ‘Unhandled’ treatment. This allowed us to balance our comparisons with respect to time of sampling and any other environmental changes in the sampling area that might influence humidity production.

### Robot sampling procedure

Humidity within sample headspace (the headspaces of flower, leaf and tube control samples) was measured using a modified version of the robot humidity transect method used in Harrap et al. (2020, 2021); unless stated otherwise, all aspects of the humidity transects conducted are as described in those publications.

Humidity sampling was conducted by a Staubli RX 160 robot arm (Pfäffikon, Switzerland). Here, background humidity is measured by a humidity probe (DHT-22 humidity probe, Aosong Electronics, Huangpu, China) placed within the lab space, the ‘background’ probe. Sample headspace humidity measurements are taken with an identical probe mounted to the robot arm, the ‘focal’ probe (figure 2). During sampling, the robot moves the focal probe through the headspace of a sample in a set sequence of transects, conducting first an *x* axis transect (horizontal) and then a *z* axis transect (vertical). During these two transects, the arm stops to take humidity measurements at set measurement points (figure 3). Following completion of *x* and *z* transects the arm conducted a probe calibration step, allowing measurements to account for variation in humidity estimation of the same humidity levels between probes. The positions of these transects, and humidity measurement points within them, for each sample are calculated by the robot in three-dimensional space relative to a manually input ‘transect central point’. This ‘transect central point’ for flower and tube control samples was set to a point in space 5mm above the centre of the flower or tube and 5 mm higher than its highest point. For leaf samples, a point in space above the centre of its horizontal span and 5 mm higher than the highest point on this horizontal span, was set as the transect central point (figure 3). Pairs of samples (prepared as described above) were presented to the robot arm and positions of transect central points (as well as positions required for probe calibration steps) input manually into the robot’s memory. The robot’s autonomous sequence was then activated, and the lab space was vacated. All robot sampling commenced within one hour of sample collection.

**Figure 2:**
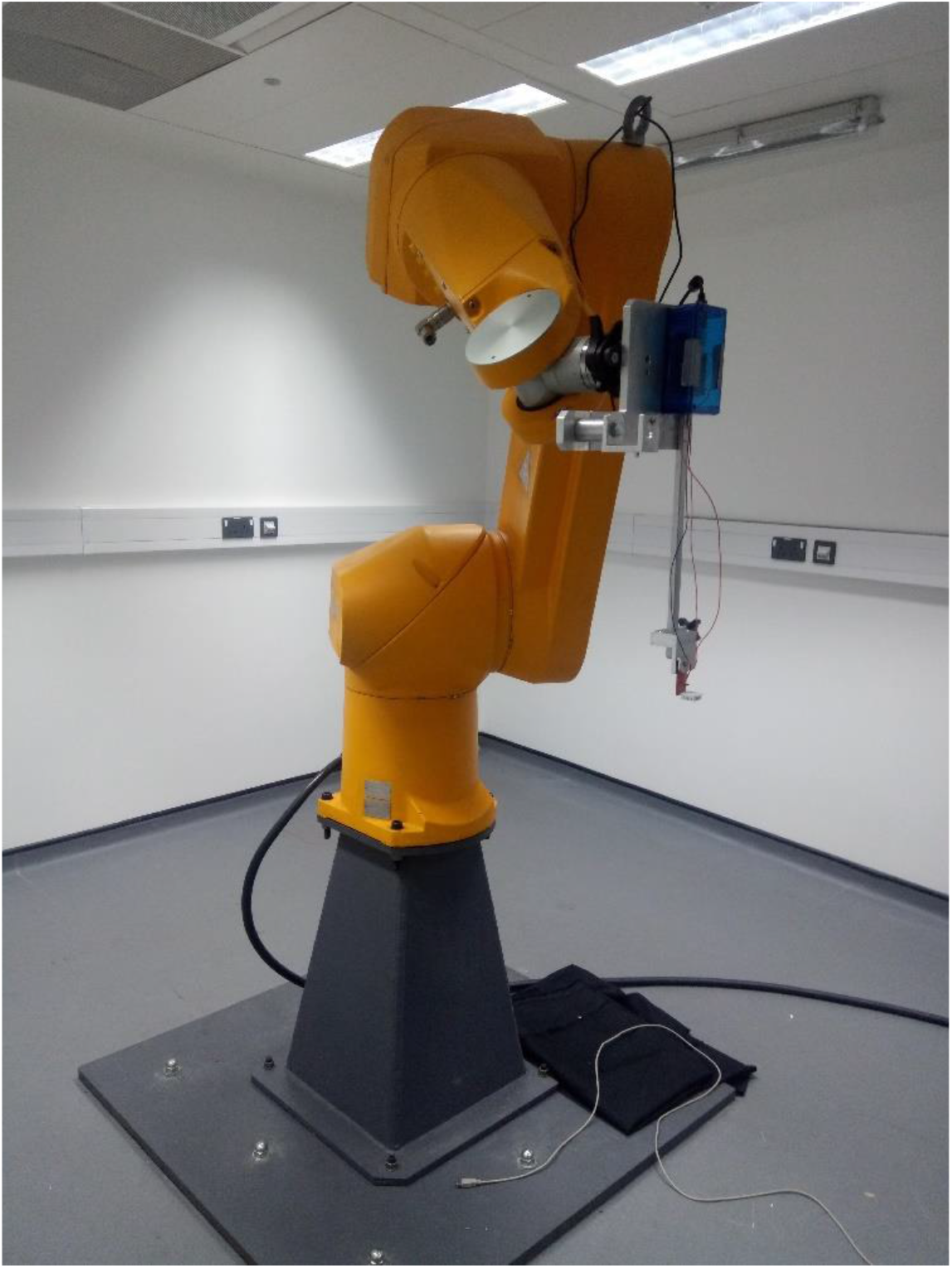
The robot arm and lab space used for headspace humidity sampling. Of note is the focal humidity probe on a 30mm bar mounted to the end of the robot arm. Samples and the background probe would be placed on a table below the robot. For further information of robot setup see Harrap et al. (2020).

**Figure 3:**
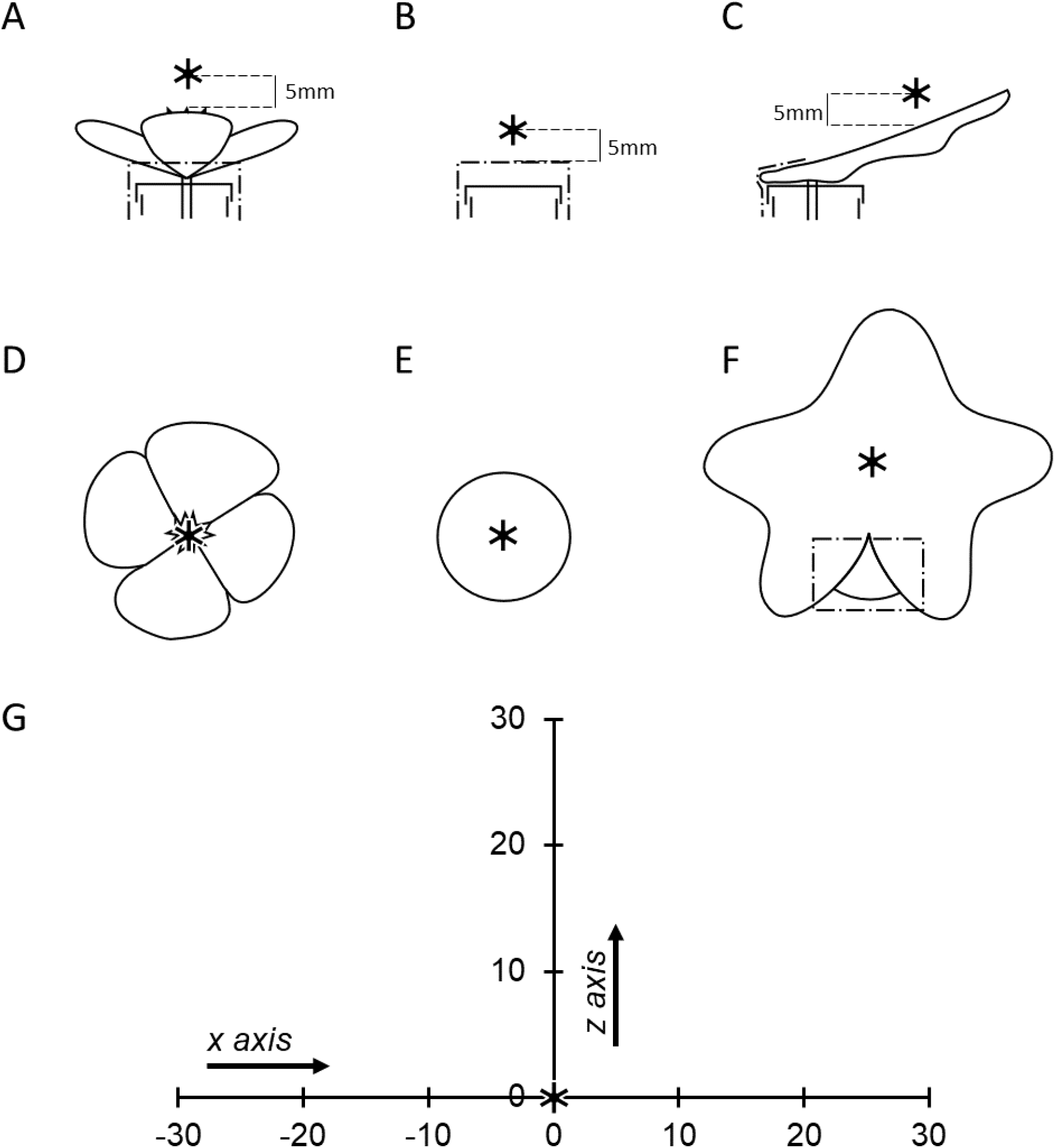
Spatial layout of humidity transects and measurement points within them relative to samples. Throughout all diagrams the ‘transect central point’ is represented by the bold asterisk ‘*’. A-C show the location of the transect central point relative to a flower (*Eschscholzia californica*), tube and leaf (*Hedera helix*) sample respectively when viewed down the horizontal *x* axis transect. D-F show the location of the transect central point relative to a flower, tube and leaf sample respectively when viewed down the vertical *z* axis transect. In each of these diagrams the robot would move towards the viewer while conducting the respective transects. Dot-dash lines indicate paper and tape supports of samples. G shows the spatial layout of the humidity headspace measured above a sample, when viewed in cross section sideways on. Each measurement point is marked with a dash and described by the offset distance along that transect (in millimetres) relative to the transect central point (*x* = 0 and *z* = 0). Arrows indicate the directions the probe travels during each transect.

Upon activation, the arm randomly chose a sample from within a pair and then conducted *x* and then *z* transects, measuring humidity at measurement points throughout (figure 3), followed by a probe calibration step. This same sequence of transects and calibration was then repeated for the other sample in the pair. After measuring both paired samples, the arm repeated this sequence on both samples in the same sampling order. Robot sampling was conducted between 2019/08/08 to 2019/08/23 and between 2020/07/06 to 2020/09/28 with sampling occurring throughout the day (between 0900 and 2000). Further detail of the layout of the sampling area and the robot sampling procedure can be found in Harrap et al. (2020) and Supplementary Information 1.

The arm stopped for 230 seconds at each measurement point on both transects (figure 3). The first 30 seconds of this was settling time to account for small disruptions from arm motion (see Harrap et al 2020). The following 200 seconds was the ‘measurement period’ for that measurement point on that transect. During the measurement period the focal probe sampled humidity continuously, and this focal humidity at the point was corrected based on the probe control step to give corrected focal humidity (*f*_*corrected*_), which accounts for variation between probes in humidity estimation of the same humidity levels (see Harrap et al., 2020 and Supplementary Information 1 for further detail). Simultaneously, the background probe measured the background humidity (*f*_*background*_). During 200 seconds, each probe could sample humidity approximately 100 times. Change in humidity, ΔRH, between the focal and background probe was then calculated as

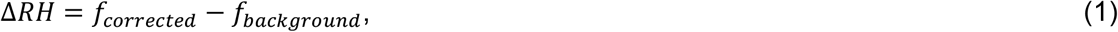

where a positive value of *ΔRH* indicates the focal probe has detected an increase in humidity in the flower’s headspace relative to the background humidity. This same measurement procedure of 30 seconds waiting time and 200 seconds measurement was also carried out during probe control measurements.

Although similar, the humidity sampling procedure used here differs in several ways from that in Harrap et al. (2020, 2021). Here, we reduced the number of measurement points on the transects, the number of replicate transects the arm conducted on each individual sample, and the number of samples presented to the arm at a time. This was done to reduce the length of each transect sequence to approximately 2 hours from robot activation to completion of sampling on the last replicate transect of a sample pair (previously 21 hours). Cutting a plant organ can separate its hormonal controls and interfere with water uptake (Huang et al., 2018; Lü et al., 2011; van Doorn, 1997), as can antitranspirant treatments (Davenport et al., 1972; Neumann, 1974), leading to drying out or wilting and changes in transpiration activity, even without blocking by antitranspirants. These effects can take time to develop, with cut flowers showing normal cycles of transpiration and water uptake within the timescales of the current or previous robot sampling procedure (Huang et al., 2018; Lü et al., 2011). Regardless, the shorter sampling sequence ensured that separated flowers were fresh, were not wilting, and were functioning more normally in terms of water uptake and transpiration where treatments were not applied.

### Analysis of humidity production

Change in humidity, *ΔRH*, as measured by the robot sampling procedure, has been found to be highly consistent within each measurement period, the *c*.100 measurements made at each sampling point, on each replicate, on each sample (Harrap et al., 2020). We therefore used the calculated the mean *ΔRH* for each measurement point and used these in our analyses.

Analysis of sample humidity production involved two steps. In the first part, the best fitting structure of humidity for each sample type was found, using models of humidity structure that allowed humidity production to vary as required with treatments. For each sample type a series of linear models were fitted to the data of the *x* and *z* axis humidity transects. These different models described different humidity structures and allowed humidity to vary with treatment. Sample identity (the identity of the individual flower, leaf or tube) was included in all models, as a random factor influencing humidity intensity (model intercept). The most complex models allowed humidity to vary with replicate transects and show a quadratic structure in the *x* axis and a logarithmic structure in the *z* axis. All other models were simplified versions of these. How well these different models described humidity structure was compared using AIC to identify the best-fitting model of *x* and *z* axis humidity structure for each sample type, using the model section criteria described in Richards (2008). These models and the associated analysis are comparable to those applied to data collected previously with this method (Harrap et al., 2020, 2021), with the additional treatment effects included in the models. The structure of these models is described in Supplementary Information 2.

In the second part of the analysis the effects of treatment on humidity production in each sample type were evaluated. Here versions of the best fitting model of humidity structure with different treatment effects were fitted to the data of each sample type. These models differed in how humidity production changed with treatments, treatment had either: no effect on humidity production, the *T0* (*x* axis) and *Tz0* (*z* axis) models; an effect on humidity intensity only (model intercept), the *T1* and *Tz1* models; an effect on humidity structure only, the *T2* and *Tz2* models; an effect on both humidity intensity and structure the *T3* and *Tz3* models; or an effect on both humidity intensity and structure as well as how humidity changes with replicate effects, the *T4* and *Tz4* models. Where the best fitting model of humidity structure for a sample, as selected above, did not include changes in humidity with replicate effects the *T4* and *Tz4* models were not fitted to the data (as there are no replicate effects for treatment to alter). Depending on the best performing humidity structure model, either the *T3* and *Tz3* or the *T4* and *Tz4* represent the ‘full’ model selected in the humidity structure step (it was possible, if a flat model with no replicate effects was favoured, for *T1* and *Tz1* be the ‘full’ model at this stage, this was not the case in any of our samples).

In both tube and leaf samples there were only two treatments (‘Untreated’ and ‘Gel’), so the models described up to this point identify the nature of treatment’s effects. However, in flower samples there were three treatments (‘Unhandled’, ‘Untreated’ and ‘Gel’). To further assess the effects of these three different treatments additional variants of the treatment effects models described above were fitted to the data that grouped together the effects of different treatments. These were identified using a subscript after the model names described above: a lack of a subscript entry indicating all treatments differ; ‘TCwP’, Untreated and Gel treatments are grouped together; ‘TCwU’, Untreated and Unhandled treatments are grouped together; ‘TPwU’, Gel and Unhandled treatments are grouped together. For example, model *T3*_*TCwP*_ describes treatment effects on both the humidity intensity and structure, but groups the ‘Gel’ and ‘Untreated’ treatments together, thus only the ‘Unhandled’ treatment differs. The *T0* and *Tz0* models describe no treatment effects, thus there were no further variants of these models. The structure of treatment effect models is described in Supplementary Information 2.

How well these various treatment effects models described humidity structure was compared using AIC to identify the best-fitting treatment effects in *x* and *z* axis for each sample type, using criteria described in Richards (2008). This two-step process of selecting a humidity structure model and then a treatment effect model was favoured as it reduced the complexity of the analysis, reducing the number of models fit to the data. Following identification of the model that best described treatment effects of each sample, humidity intensity summary values 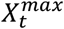 (the position of the mean peak in humidity production over the x axis transect, relative to the transect central point) and 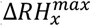 (the average peak in humidity production over the *x* axis transect) were calculated according to the best fitting treatment model for each treatment of each sample and each replicate transect when it was included in the best fitting model. These summary values, respectively, give a conservative estimate of humidity structure symmetry and humidity intensity produced by each sample and treatment. For further detail on summary values and their calculation see Harrap et al. (2020) and Supplementary Information 3.

### Monitoring of sample temperature

Floral transpiration can play an important role in floral temperature regulation (Patiño & Grace, 2002) so blocking transpiration may influence floral temperature. Understanding antitranspirant treatments’ influence on floral temperature may help explain floral humidity changes and the relationship between these traits. Temperature was therefore monitored alongside humidity sampling for a subset of samples (table 1). In these instances, a thermal camera (FLIR *E60bx*, FLIR systems Inc., Wilsonville, USA) was mounted on a tripod viewing the sample pair from an elevated side on angle. A small (*c*. 10 x 20cm) aluminium foil multidirectional mirror was placed in view of the camera against with the bottom of the horticultural tube rack. This fully charged thermal camera was automated to take a thermal image upon activation just before humidity sampling began and every subsequent 15 minutes using the FLIR Tools software live feed functionality (FLIR systems INC, 2015) and autoclicker software (written within AutoHotkey, Mallett & AutoIt Team, 2014) running on an attached laptop placed within the sampling area. Neither the thermal camera nor the laptop generated considerable heat or had fan components that might influence turbulence within the sampling area.

The capture of a thermal images during sampling continued until the camera ran out of battery or the camera-PC connection was lost (which could be identified by sequential duplicate images that did not show robot motion which should have been visible). Due to constraints of the software, loss of the PC connection often occurred before the end of floral sampling. Thermal images captured more than 151 minutes after the start of sampling were also discarded as sampling had finished by this point. All remaining thermal images were included in our analysis of floral temperature over the sampling period, regardless of when camera connection was lost.

The temperature of samples was taken from the thermal images in FLIR tools using a manually placed point measurement at the centre of the visible portion of each flower or control tube. During thermographic measurements, target emissivity was set to 0.98, a value appropriate for floral and vegetative tissue (Harrap & Rands, 2021b), and reflected temperature measured using the multidirectional mirror placed in frame (Harrap et al., 2018): distance was assumed to be 1m, while humidity and environmental temperature, which have only minor effects on measurements (Usamentiaga et al., 2014; Vollmer & Möllmann, 2017), were set to 50% and 20°C respectively. Application of gel, being composed of different material to plant tissue and differing in texture, may alter object emissivity. Although similar Vaseline mixtures have been found to lower the infrared emissivity of targets like skin, this effect is very small (Steketee, 1976) and previous thermography work measuring Vaseline-treated leaves with thermographic tools deemed it unnecessary to change emissivity settings between treated and untreated leaves (Jones, 1999; Leinonen & Jones, 2004; Grant et al., 2006; Stoll & Jones, 2007). Consequently, we chose to not change emissivity for gel-covered targets.

As different treatments were applied to flower species (‘Unhandled’, ‘Untreated’ and ‘Gel’), and the Dry Tube controls (‘Untreated’ and ‘Gel’), the temperatures of each flower species and the Dry tube controls were analysed separately to avoid rank deficiencies and avoiding complex three factor interactions. For both species and the Dry Tube controls the effects of treatment and time elapsed during sampling (measured as decimalized minutes) was assessed using a repeated measures ANOVA, including both flower or control identity and the pairings of samples (i.e. which samples were monitored at the same time) as a random factors.

## RESULTS

### Control samples

In the Dry Tube control, we saw little change in the humidity during sampling (figure 4A and B, table 2). Best fitting models, according to AIC (table 2), indicated application of gel had no influence on humidity production in either *x* or *z* transects. This led to models without treatment effects (*T0* and *Tz0*) being favoured, which indicates the presence of gel on its own did not produce or reduce the humidity of Dry Tube controls.

**Table 2:**
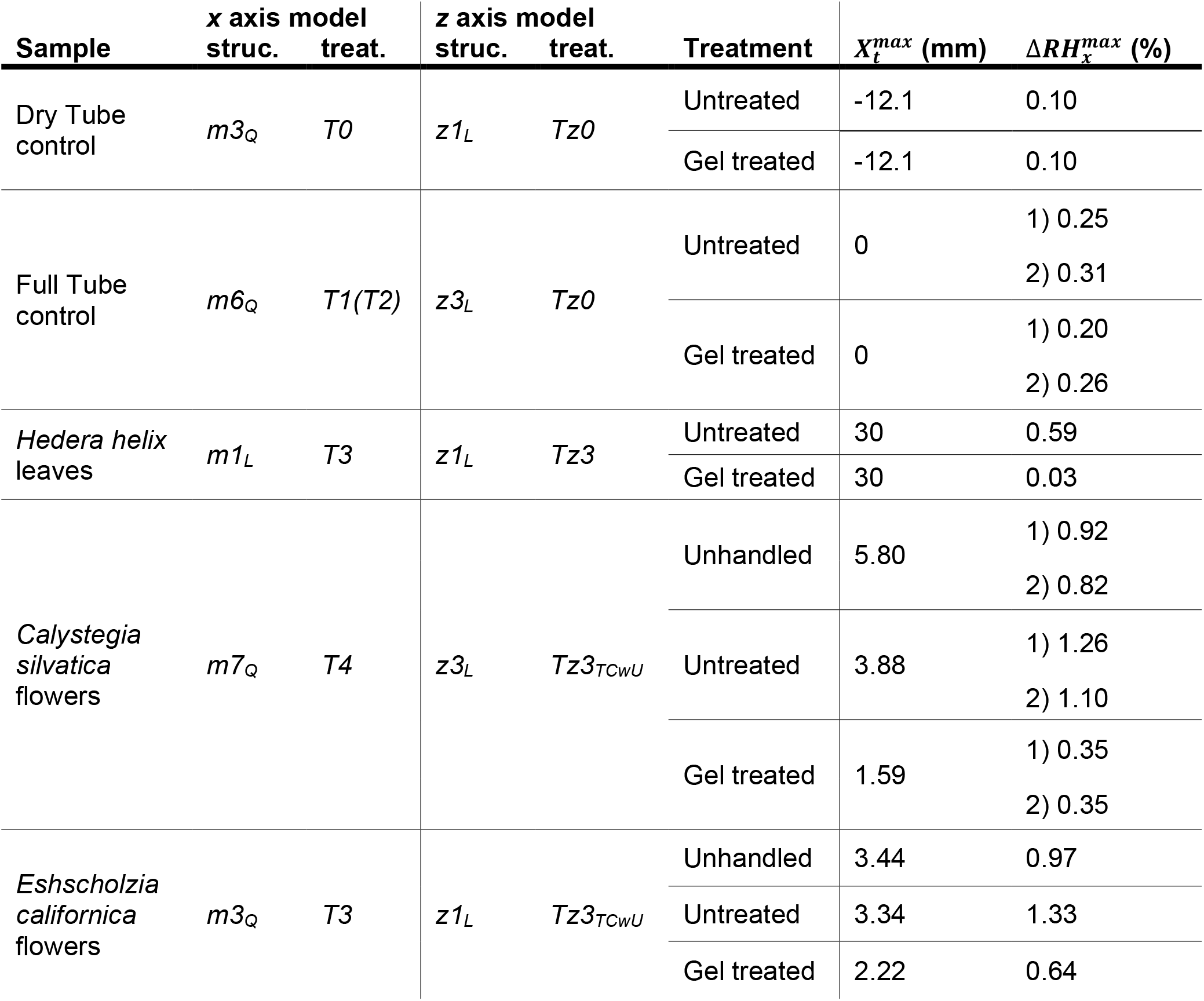
Summary of humidity and the influences of treatments on control samples, *Hedra helix* leaves and flowers (hereafter ‘samples’). For each transect (*x* and *z*) of each sample the best fitting model of humidity structure (column ‘struc.’) and treatment effects (column ‘treat.’) are given. Where more than one comparable models are best both are given, model outside bracket indicates the lower AIC model used for subsequent calculations. Subscript letters following ‘struc.’ models indicate the shape of humidity described by the best fitting model: ‘*L*’ a linear (*x* axis) or log-linear (*z* axis) structure, and ‘*Q*’ a quadratic structure (*x* axis only). For each treatment of each sample 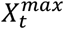 and 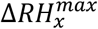 values estimated by the best fitting *x* axis ‘treat.’ model a given. Changes in humidity structure and intensity between treatments are indicated by corresponding differences in 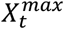 and 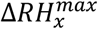 respectively. Where the best model of a sample indicates a change in 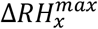 with replicate transects 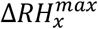 of both are given: ‘1)’ indicates the first transects; ‘2)’ the second. For further detail on model identity and structure see Supplementary Information 2. For expanded results of AIC tests for each sample type see Supplementary Information 4.

**Figure 4:**
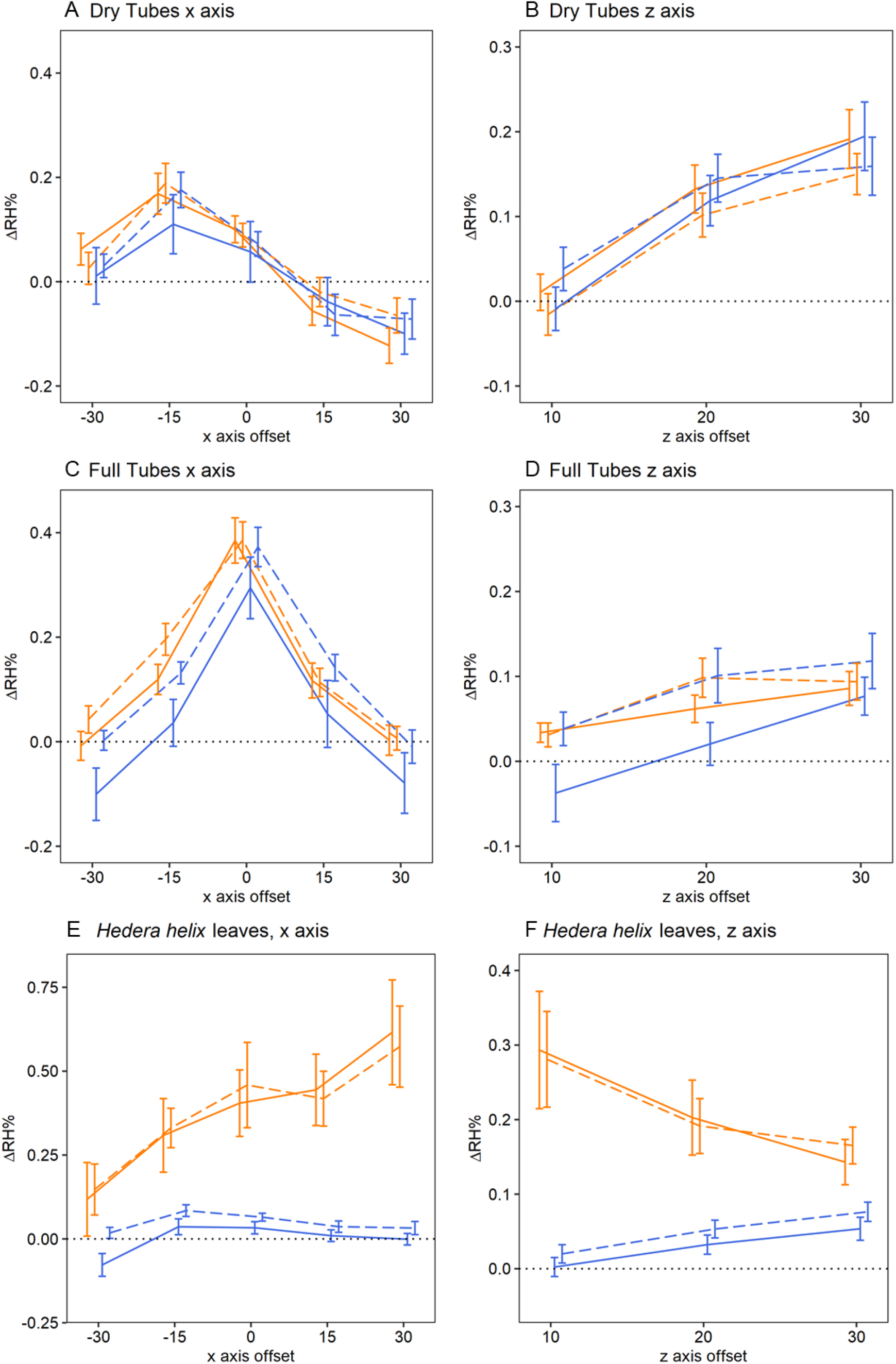
The effect of antitranspirant gel treatments on the humidity production of tube controls and leaf samples. Plots show mean difference in humidity relative to the background (ΔRH) across the *x* (A, C, E) and *z* (B, D, F) axis transects of: A, B, Dry tube controls; C, D, Full tube controls; and E, F, *Hedera helix* leaves. All axis offsets are relative to the transect central point and in millimetres. The thin dotted line indicates a 0% change in humidity (the background level). Bold lines plot the mean ΔRH of each treatment of each sample at each replicate transect. Error bars represent ±S.E.M. Colour indicates treatment: orange the ‘Untreated’ treatment; blue the ‘Gel’ treatment. Dashing of lines indicates the transect replicate: solid, first transect; dashed, the second transect. Note that positioning of lines and bars is offset from the measurement point in the *x* and *z* axis for clarity.

When horticultural tubes were filled with water (the Full Tube control), humidity production increased, as seen by the slightly elevated 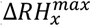 values (table 2). According to the best fitting humidity structure model, humidity production showed a quadratic structure with peak humidity at the centre of the transect (figure 4C), indicating the source of this humidity was evaporation of water through the hole in tube lids. A small rise in humidity with replicate transects suggests that this evaporation slowly accumulated over the sampling period after tubes were moved to their position in the sampling area. Application of gel to the Full Tube control influenced humidity production slightly. This small effect of treatment meant treatment effect models that allowed gel treatment to either cause a 0.05% drop in relative humidity (*T1* treatment model) or a small change in humidity structure (*T2* treatment model) performed best but were comparable in terms of AIC (table 2, supplementary information 4). As gel treatment did not change humidity production in the Dry Tube control, this small influence of gel treatment is likely due to gel slightly blocking evaporation, pooling in such a way that it obscures or effectively narrows the hole in the tube’s lid. A similar small effect of obscuring the hole in the tubes lid was observed in Harrap et al. (2020).

In the *z* axis transect, both control samples showed a (log-)linear structure according to the best fitting humidity structure model (figure 4B and D). Humidity production, Δ*RH*, nearer the control samples (*z* offset = 10mm) was approximately 0%, with humidity rising by small amounts (*c*.0.1-0.2%) with increased distance. Treatments had no effect on humidity production in the *z* axis transect in either control, leading to models with no treatment effects (*Tz0* models) being favoured in both control samples (table 2).

Best-fitting models of the Full Tube control suggest extraneous humidity sources were low (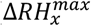 was between 0.25 and 0.31%, that of the Untreated Full tube control). However, leaf and flower samples also block the hole in the horticultural tube lid. Consequently, extraneous humidity sources are likely to be lowered in a similar manner as seen in Harrap et al. (2020) and the ‘Gel’ treated Full Tube control.

### Leaf samples

‘Untreated’ *H. helix* leaf transects found humidity about the leaf to be elevated by a small amount 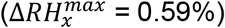. Humidity produced by untreated leaves increased across the *x* axis, and best-fitting models of leaf *x* axis transects indicated a linear humidity structure of untreated leaves (figure 4E, table 2). In *z* axis transects, humidity declined with increased distance from untreated leaves (figure 4F, table 2).

Gel application on *H. helix* leaves changed both the structure and amount of humidity produced by leaf samples, resulting in *T3* and *Tz3* treatment models being favoured (figure 4E and F, table 2). Gel treatment reduced the amount of humidity produced by leaves in the *x* axis almost completely 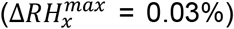. This was paired with a flattening in humidity structure over the *x* axis transect, meaning Δ*RH* remained at near zero levels across the *x* transects of treated leaves (figure 4E). In the *z* transect, humidity intensity was likewise reduced, but structure changed so that humidity rose slightly over the *z* transect (figure 4F), as in control samples. As leaf humidity generation would be predominantly through transpiration (Vezza et al., 2006; von Arx et al., 2012), the almost complete removal of leaf headspace humidity with gel treatment demonstrates the gel effectively blocks transpiration from treated tissues and consequently removes transpiration’s contribution to sample humidity.

### Flower samples

Floral humidity detected in the headspaces of ‘Unhandled’ *C. silvatica* and *E. californica* flowers (figure 5) were broadly consistent with floral humidity observed previously (Harrap et al., 2020). Best-fitting models indicated floral humidity in both species showed a quadratic structure across the *x* axis transect (figure 5A and C, table 2). In both species, 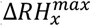 was close to 1% in Unhandled flowers. In the *z* axis transect, both species slowed a decrease in humidity with increased distance from the flower (figure 5B and D). Furthermore, best-fitting models indicated *C. silvatica* floral humidity decreased slightly with replicate transects in the x axis. In *E. californica*, humidity was not found to differ between replicate transects.

**Figure 5:**
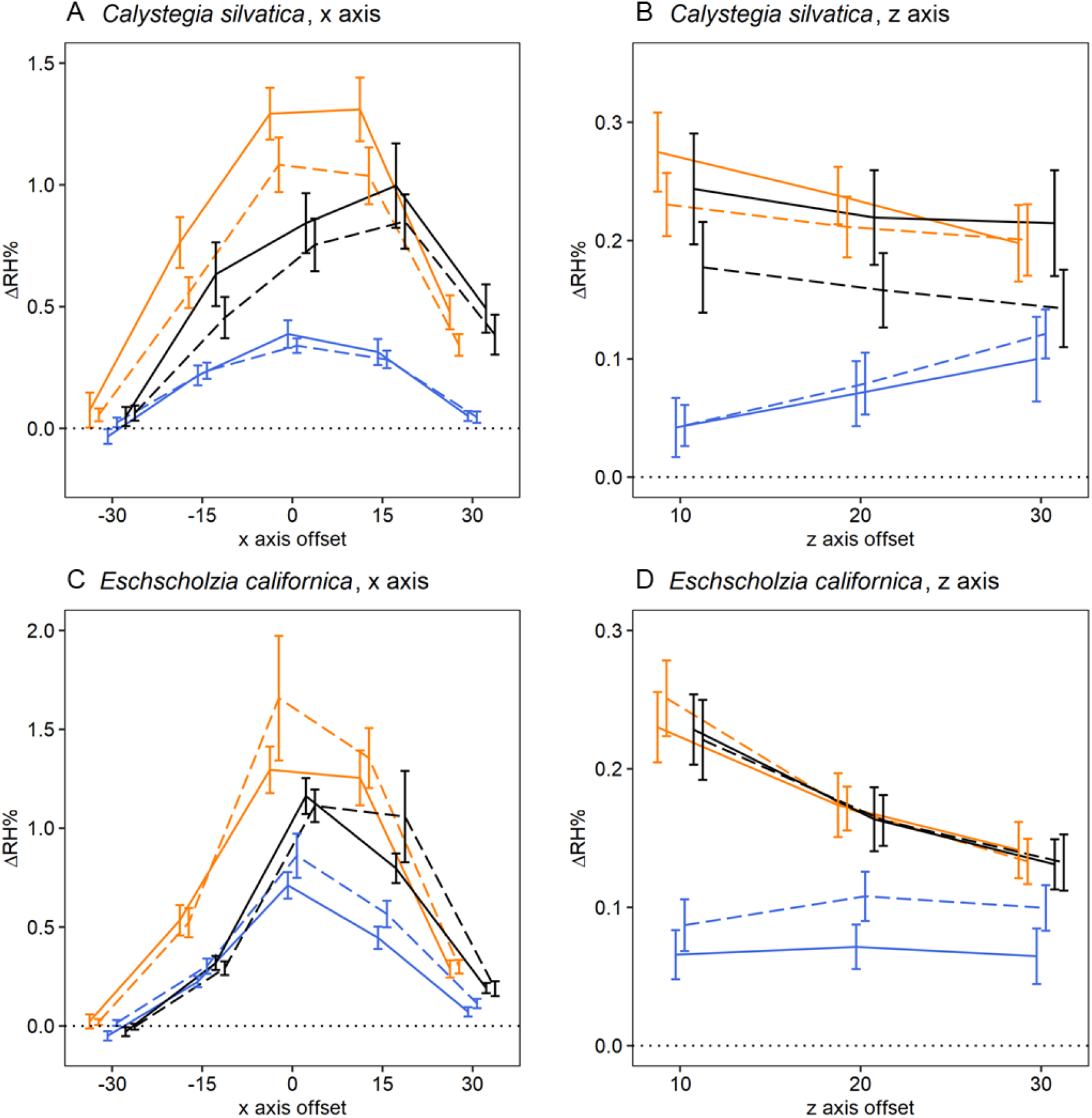
The effect of antitranspirant gel treatments on the humidity production of flowers. Plots show mean difference in humidity relative to the background (ΔRH) across the *x* (A, C) and *z* (B, D) axis transects of: A, B, *Calystegia silvatica*; C, D, *Eschscholzia californica*. Details as for figure 4, noting that colour here indicates treatment: black, the ‘Unhandled’ treatment; orange, the ‘Untreated’ treatment; blue, the ‘Gel’ treatment.

The treatments influenced both the intensity and structure of floral humidity in both transects of both species (figure 5, table 2), and influenced how *C. silvatica* humidity intensity changed with replicate transects within the *x* axis. For the *x* and *z* axis transects respectively, the *T3* and *Tz3*_*TCwU*_ treatment effect models were favoured by AIC for *E. californica*, and the *T4* and *Tz3*_*TCwU*_ models for *C. silvatica* (table 2).

Handling flowers as if gel were being applied (the ‘Untreated’ treatment) led to a small increase in humidity production compared to flowers of the same species that had not been handled (the ‘Unhandled’ treatment, figure 5). This increase in humidity production was similar between both species with 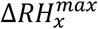 increasing in relative humidity in ‘Untreated’ flowers compared to ‘Unhandled’ flowers by 0.36% in *E. californica* and 0.34% in the initial transect of *C. silvatica*, 0.28% in the second transect. In *z* axis transects of both species, humidity was found not to differ between ‘Unhandled’ and ‘Untreated’ treatments (table 2).

In both *C. silvatica* and *E. californica*, application of gel to petal surfaces to block transpiration (the ‘Gel’ treatment) reduced floral humidity production compared to other treatments (figure 5). Humidity intensity of ‘Gel’ treated *C. silvatica* flowers was approximately a third of that in ‘Untreated’ flowers (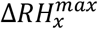 being 0.35% in both transects). In *E. californica* floral humidity intensity of ‘Gel’ treated flowers was approximately half that of ‘Untreated’ flowers (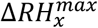 being 0.64%). These decreases were accompanied by a flattening of the *x* axis humidity structure in both species (figure 5). Additionally, in *C. silvatica* floral humidity production of gel treated flowers was more consistent, remaining at a reduced level between replicate transects. In the *z* axis transects, humidity was likewise reduced by gel treatments in both species, but in *C. silvatica* humidity was further affected with *z* axis humidity structure changing so that humidity rose slightly over the *z* transect (figure 5), much like that of Dry Tube control samples.

### Treatments and temperature of flowers and controls

Temperature monitoring of Dry Tube controls during humidity sampling revealed application of gel had no detectable effect on tube surface temperature (*minutes elapsed and gel treatment interaction effects* ANOVA, *F*_1,37_ = 0.438, *p* = 0.512; *gel treatment* ANOVA, *F*_1,37_ = 0.273 *p* = 0.605). Furthermore, tubes did not show a change in temperature over time (*minutes elapsed* ANOVA, *F*_1,37_ = 0.502, *p* = 0.483). That gel treated tubes did not differ in temperature suggests the gel itself does not generate heat. Gel treatment likewise had no effect on floral surface temperature of *E. californica* flowers (*minutes elapsed and gel treatment interaction effects* ANOVA, *F*_2,148_ = 1.023, *p* = 0.362; *gel treatment* ANOVA, *F*_2,14_ = 1.048 *p* = 0.376). *E. californica* flowers began sampling at approximately the temperature of the sampling room (22.58°C, according to ANOVA model mean estimates) and cooled slightly (at a rate of −8.5×10^−4^°C/min) during humidity sampling (*minutes elapsed* ANOVA, *F*_1,149_ = 4.716, *p* = 0.031). However, gel treatment did affect the temperature of *C. silvatica* flowers, which showed a similar change in temperature with time regardless of treatment (*minutes elapsed and gel treatment interaction effects* ANOVA, *F*_2,176_ = 1.287, p = 0.279), with flowers cooling gradually (at a rate of −6.1×10^−4^°C/min) over time (*minutes elapsed* ANOVA, *F*_1,176_ = 13.889, *p* < 0.001). Nevertheless, gel treatment had a small but significant effect on the overall temperature of the flowers (*gel treatment* ANOVA, *F*_2,16_ = 6.753, *p* = 0.008). *Post hoc* Tukey tests revealed this was due to ‘Gel’ treated flowers being, on average, 0.3°C hotter than ‘Untreated’ flowers (*Untreated-Gel comparison*, Tukey test, *p* = 0.025), ‘Untreated’ flowers again beginning sampling at approximately room temperature (22.53°C, according to ANOVA model mean estimates). No significant temperature differences were found between other treatment group pairings of *C. silvatica* flowers (*Untreated-Unhandled comparison*, Tukey test, *p* = 0.231; *Gel-Unhandled comparison*, Tukey test, *p* = 0.996).

## DISCUSSION

Antitranspirant treatment of flower petals and sepals and the resulting effects on floral humidity confirm that transpiration contributes to the generation of floral humidity. Gel-treated flowers produced less floral humidity than flowers that received no gel treatments (figure 5, table 2). This gel antitranspirant did not produce or absorb water vapour itself, confirmed by the lack of treatment effects on Dry Tube control samples (figure 4A and B), and was confirmed to block transpiration from treated plant tissues, and remove transpiration’s contribution to sample headspace humidity, shown by the removal of humidity from headspaces of gel treated leaves (figure 4D and E). Handling during gel application affected humidity, evidenced by the slightly increased floral humidity of ‘Untreated’ flowers of both species compared to ‘Unhandled’ flowers (table 2). Such changes in humidity production due to handling alone may be due to micro-abrasions and cell damage to petal surfaces during handling, or transfer of grease or moisture from fingers. Regardless, this conflating handling effect was small. Considering the results together, the reduction in humidity production of flowers with gel treatments appears to represent the effect of gel removing the contribution of transpiration from floral humidity production. Thus, the results of floral treatments confirm previous, but until now untested, inferences that floral humidity is produced in part by floral transpiration (Harrap et al., 2020; von Arx et al., 2012).

The gel did not itself generate heat, as temperatures of Dry tube controls were unaffected by treatment. Likewise, temperature of *E. californica* flowers was not influenced by gel treatments. However, gel treatment did slightly increase *C. silvatica* floral temperature. That ‘Untreated’ flowers were either the same temperature as, or slightly cooler than, gel treated flowers confirms that floral humidity seen in ‘Untreated’ flowers is not the result of elevated temperatures enhancing nectar evaporation, or transpiration, relative to gel-treated flowers. The rise in temperature with gel treatment *C. silvatica* appears due to the gel blocking transpiration, compromising the flowers’ capacity to lose heat, observed with similar antitranspirant treatments of other Convolvulaceae (Patiño & Grace, 2002). This effect may be greater in *C. silvatica* due to its larger size, limiting passive heat loss with air conduction, explaining why temperature differences were not detected in *E. californica*. That preventing transpiration impacts both temperature and humidity in *C. silvatica* provides evidence that a relationship between floral humidity and floral temperature exists though the influence of floral transpiration. That only small increases in temperature were created when heat loss from transpiration is removed suggests that flowers were not under much heat stress during sampling, thus only small amounts of heat accumulate by preventing transpirational heat loss.

Our results indicate transpiration can have quite large contributions to floral humidity generation and species can vary in the extent that floral humidity is produced by transpiration. Although total floral humidity production in Untreated flowers was similar in both species, transpiration was found to have a greater contribution to floral humidity production in *C. silvatica*, accounting for a larger part of the total floral humidity produced than in *E. californica*. Furthermore, in *C. silvatica* it appears transpiration has a greater contribution relative to other sources. The differences between gel-treated flowers and flowers in other treatments indicates transpiration accounts for about a half of the total floral humidity produced by *E. californica*, and two thirds of that produced by *C. silvatica*. However, floral transpiration may have a greater contribution to total floral humidity then indicated by our treatments. Neither floral reproductive structures nor areas proximal to them (including nectaries) were gel treated. These structures may also transpire and this will contribute to the remaining humidity production of gel-treated flowers. Additionally, the elevated humidity detected in the floral headspace of gel-treated flowers still includes humidity from sources extraneous to the flower. These ‘extraneous sources’ may account for differences between focal and background probes of up to 0.31% relative humidity (that of the ‘Untreated’ full tube).

Several influences determine the capacity of a flower to generate floral humidity (Harrap et al., 2020). These include evaporation of nectar, which produces humidity, and floral structure, which influences how it accumulates in flower headspaces (von Arx et al., 2012). Our results confirm, as previously inferred, that transpiration also contributes to the amount of floral humidity generated. This means traits that influence floral transpiration will impact the capacity of flower species to produce floral humidity, and differences in these traits may help explain the diversity of floral humidity produced across the angiosperms (Harrap et al., 2020).

## SUPPLEMENTARY DATA

### Supplementary information

This supplementary information file contains the following materials as four sections: Supplementary information 1, Extended detail of robot sampling procedure; Supplementary information 2, Further detail of models and model simplification procedures; Supplementary information 3, Summary value calculation; and Supplementary information 4, AIC tables.

## SUPPLEMENTARY MATERIALS

One file containing 3 supplementary information sections (providing further extended details of methods) and 8 supplementary tables

## ACKNOWLEDGEMENTS

This research was funded by the Bristol Centre for Agricultural Innovation.

## AUTHOR CONTRIBUTIONS

SAR provided resources, supervision, project administration and acquired funding. MJMH developed the methodology for this study, conducted the investigation, data curation, visualisation and conducted formal analysis. Both authors were involved in study conceptualization and together wrote the manuscript.

## DATA AVAILABILTY STATEMENT

Datafiles and code for the analyses described in this study and figure generation are openly available on the Figshare database at https://doi.org/10.6084/m9.figshare.14350547 (Harrap & Rands, 2021a).

## REFERENCES

Ashman, T.-L., Knight, T. M., Steets, J. A., Amarasekare, P., Burd, M., Campbell, D. R., Dudash, M. R., Johnston, M. O., Mazer, S. J., Mitchell, R. J., Morgan, M. T., & Wilson, W. G. (2004). POLLEN LIMITATION OF PLANT REPRODUCTION: ECOLOGICAL AND EVOLUTIONARY CAUSES AND CONSEQUENCES. Ecology, 85(9), 2408–2421. https://doi.org/10.1890/03-8024

Azad, A. K., Sawa, Y., Ishikawa, T., & Shibata, H. (2007). Temperature-dependent stomatal movement in tulip petals controls water transpiration during flower opening and closing. Annals of Applied Biology, 150(1), 81–87. https://doi.org/10.1111/j.1744-7348.2006.00111.x

Baude, M., Kunin, W. E., Boatman, N. D., Conyers, S., Davies, N., Gillespie, M. A. K., Morton, R. D., Smart, S. M., & Memmott, J. (2016). Historical nectar assessment reveals the fall and rise of floral resources in Britain. Nature, 530(7588), 85–88. https://doi.org/10.1038/nature16532

Buschhaus, C., Hager, D., & Jetter, R. (2015). Wax layers on Cosmos bipinnatus petals contribute unequally to total petal water resistance. Plant Physiology, 167(1), 80–88. https://doi.org/10.1104/pp.114.249235

Cheng, G., Huang, H., Zhou, L., He, S., Zhang, Y., & Cheng, X. (2019). Chemical composition and water permeability of the cuticular wax barrier in rose leaf and petal: A comparative investigation. Plant Physiology and Biochemistry, 135, 404–410. https://doi.org/10.1016/j.plaphy.2019.01.006

Corbet, S. A., Unwin, D. M., & Prŷs-Jones, O. E. (1979). Humidity, nectar and insect visits to flowers, with special reference to Crataegus, Tilia and Echium. Ecological Entomology, 4(1), 9–22. https://doi.org/10.1111/j.1365-2311.1979.tb00557.x

Corbet, S. A., Willmer, P. G., Beament, J. W. L., Unwin, D. M., & Prŷs-Jones, O. E. (1979). Post-secretory determinants of sugar concentration in nectar. Plant, Cell & Environment, 2(4), 293–308. https://doi.org/10.1111/j.1365-3040.1979.tb00084.x

Davenport, D. C., Fisher, M. A., & Hagan, R. M. (1972). Some counteractive effects of antitranspirants. Plant Physiology, 49(5), 722–724.

FLIR systems INC. (2015). FLIR tools (PENDING) [Computer software]. FLIR systems INC.

Gates, D. M. (1968). Transpiration and leaf temperature. Annual Review of Plant Physiology, 19(1), 211–238. https://doi.org/10.1146/annurev.pp.19.060168.001235

Grant, O. M., Chaves, M. M., & Jones, H. G. (2006). Optimizing thermal imaging as a technique for detecting stomatal closure induced by drought stress under greenhouse conditions. Physiologia Plantarum, 127(3), 507–518. https://doi.org/10.1111/j.1399-3054.2006.00686.x

Guo, Y., Busta, L., & Jetter, R. (2017). Cuticular wax coverage and composition differ among organs of Taraxacum officinale. Plant Physiology and Biochemistry, 115, 372–379. https://doi.org/10.1016/j.plaphy.2017.04.004

Hajibagheri, M. A., Hall, J. L., & Flowers, T. J. (1983). The structure of the cuticle in relation to cuticular transpiration in leaves of the halophyte Suaeda maritima (L.) Dum. New Phytologist, 94(1), 125–131. https://doi.org/10.1111/j.1469-8137.1983.tb02728.x

Harrap, M. J. M., de Ibarra, N. H., Knowles, H. D., Whitney, H. M., & Rands, S. A. (2021). Bumblebees can detect floral humidity. BioRxiv. https://doi.org/10.1101/2021.03.19.436119

Harrap, M. J. M., Hempel de Ibarra, N., Knowles, H. D., Whitney, H. M., & Rands, S. A. (2020). Floral humidity in flowering plants: A preliminary survey. Frontiers in Plant Science, 11, 249. https://doi.org/10.3389/fpls.2020.00249

Harrap, M. J. M., Hempel de Ibarra, N., Whitney, H. M., & Rands, S. A. (2018). Reporting of thermography parameters in biology: A systematic review of thermal imaging literature. Royal Society Open Science, 5, 181281. https://doi.org/10.1098/rsos.181281

Harrap, M. J. M., & Rands, S. A. (2021a). Data from “The role of transpiration in floral humidity generation.” Figshare Database. https://doi.org/10.6084/m9.figshare.14350547

Harrap, M. J. M., & Rands, S. A. (2021b). Floral infrared emissivity estimates using simple tools. Plant Methods, 17(1), 23. https://doi.org/10.1186/s13007-021-00721-ws

Hase, A. V., Cowling, R. M., & Ellis, A. G. (2006). Petal movement in cape wildflowers protects pollen from exposure to moisture. Plant Ecology, 184(1), 75–87. https://doi.org/10.1007/s11258-005-9053-8

Hew, C. S., Lee, G. L., & Wong, S. C. (1980). Occurrence of non-functional stomata in the flowers of tropical orchids. Annals of Botany, 46(2), 195–201. https://doi.org/10.1093/oxfordjournals.aob.a085907

Hicks, D. M., Ouvrard, P., Baldock, K. C. R., Baude, M., Goddard, M. A., Kunin, W. E., Mitschunas, N., Memmott, J., Morse, H., Nikolitsi, M., Osgathorpe, L. M., Potts, S. G., Robertson, K. M., Scott, A. V., Sinclair, F., Westbury, D. B., & Stone, G. N. (2016). Food for pollinators: Quantifying the nectar and pollen resources of urban flower meadows. PLoS One, 11(6), e0158117. https://doi.org/10.1371/journal.pone.0158117

Huang, X., Lin, S., He, S., Lin, X., Liu, J., Chen, R., & Li, H. (2018). Characterization of stomata on floral organs and scapes of cut ‘Real’ gerberas and their involvement in postharvest water loss. Postharvest Biology and Technology, 142, 39–45. https://doi.org/10.1016/j.postharvbio.2018.04.001

Jolliet, O., & Bailey, B. J. (1992). The effect of climate on tomato transpiration in greenhouses: Measurements and models comparison. Agricultural and Forest Meteorology, 58(1), 43–62. https://doi.org/10.1016/0168-1923(92)90110-P

Jones, H. G. (1999). Use of thermography for quantitative studies of spatial and temporal variation of stomatal conductance over leaf surfaces. Plant, Cell & Environment, 22(9), 1043–1055. https://doi.org/10.1046/j.1365-3040.1999.00468.x

Leinonen, I., & Jones, H. G. (2004). Combining thermal and visible imagery for estimating canopy temperature and identifying plant stress. Journal of Experimental Botany, 55(401), 1423–1431. https://doi.org/10.1093/jxb/erh146

Leonard, A. S., Dornhaus, A., & Papaj, D. R. (2011). Forget-me-not: Complex floral displays, inter-signal interactions, and pollinator cognition. Current Zoology, 57(2), 215–224. https://doi.org/10.1093/czoolo/57.2.215

Leonard, A. S., Dornhaus, A., & Papaj, D. R. (2012). Why are floral signals complex? An outline of functional hypotheses. In S. Patiny (Ed.), Evolution of plant–pollinator relationships (pp. 279–300). Cambridge University Press. dx.doi.org/10.1017/CBO9781139014113.010

Leonard, A. S., & Masek, P. (2014). Multisensory integration of colors and scents: Insights from bees and flowers. Journal of Comparative Physiology A, 200, 463–474. https://doi.org/10.1007/s00359-014-0904-4

Lü, P., Huang, X., Li, H., Liu, J., He, S., Joyce, D. C., & Zhang, Z. (2011). Continuous automatic measurement of water uptake and water loss of cut flower stems. HortScience, 46(3), 509–512. https://doi.org/10.21273/HORTSCI.46.3.509

Mallett, C., & AutoIt Team. (2014). AutoHotkey—A scripting language for desktop automation (1.133.02) [Computer software].

Nepi, M., Franchi, G. G., & Padni, E. (2001). Pollen hydration status at dispersal: Cytophysiological features and strategies. Protoplasma, 216(3), 171. https://doi.org/10.1007/BF02673869

Neumann, P. M. (1974). Senescence of Attached Bean Leaves Accelerated by Sprays of Silicone Oil Antitranspirants. Plant Physiology, 53(4), 638–640. https://doi.org/10.1104/pp.53.4.638

Nordström, K., Dahlbom, J., Pragadheesh, V. S., Ghosh, S., Olsson, A., Dyakova, O., Suresh, S. K., & Olsson, S. B. (2017). In situ modeling of multimodal floral cues attracting wild pollinators across environments. Proceedings of the National Academy of Sciences of the USA, 114(50), 13218–13223. https://doi.org/10.1073/pnas.1714414114

Patiño, S., & Grace, J. (2002). The cooling of convolvulaceous flowers in a tropical environment. Plant, Cell & Environment, 25(1), 41–51. https://doi.org/10.1046/j.0016-8025.2001.00801.x

Raguso, R. A. (2004). Flowers as sensory billboards: Progress towards an integrated understanding of floral advertisement. Current Opinion in Plant Biology, 7, 434–440. https://doi.org/10.1016/j.pbi.2004.05.010

Raine, N. E., & Chittka, L. (2008). The correlation of learning speed and natural foraging success in bumble-bees. Proceedings of the Royal Society B: Biological Sciences, 275(1636), 803–808. https://doi.org/10.1098/rspb.2007.1652

Rawson, H. M., Begg, J. E., & Woodward, R. G. (1977). The effect of atmospheric humidity on photosynthesis, transpiration and water use efficiency of leaves of several plant species. Planta, 134(1), 5–10. https://doi.org/10.1007/BF00390086

Richards, S. A. (2008). Dealing with overdispersed count data in applied ecology. Journal of Applied Ecology, 45(1), 218–227. https://doi.org/10.1111/j.1365-2664.2007.01377.x

Schiestl, F. P., & Johnson, S. D. (2013). Pollinator-mediated evolution of floral signals. Trends in Ecology & Evolution, 28(5), 307–315. https://doi.org/10.1016/j.tree.2013.01.019

Schreiber, L. (2001). Effect of temperature on cuticular transpiration of isolated cuticular membranes and leaf discs. Journal of Experimental Botany, 52(362), 1893–1900. https://doi.org/10.1093/jexbot/52.362.1893

Simon, N. M. L., Graham, C. A., Comben, N. E., Hetherington, A. M., & Dodd, A. N. (2020). The Circadian Clock Influences the Long-Term Water Use Efficiency of Arabidopsis. Plant Physiology, 183(1), 317–330. https://doi.org/10.1104/pp.20.00030

Steketee, J. (1976). The influence of cosmetics and ointments on the spectral emissivity of skin (skin temperature measurement). Physics in Medicine and Biology, 21(6), 920–930. https://doi.org/10.1088/0031-9155/21/6/002

Stoll, M., & Jones, H. G. (2007). Thermal imaging as a viable tool for monitoring plant stress. OENO One, 41(2), 77–84. https://doi.org/10.20870/oeno-one.2007.41.2.851

Usamentiaga, R., Venegas, P., Guerediaga, J., Vega, L., Molleda, J., & Bulnes, F. G. (2014). Infrared thermography for temperature measurement and non-destructive testing. Sensors, 14(7), 12305–12348. https://doi.org/10.3390/s140712305

van Doorn, W. G. (1997). Water relations of cut flowers. Horticultural Reviews, 18, 1–85.

Vezza, M., Nepi, M., Guarnieri, M., Artese, D., Rascio, N., & Pacini, E. (2006). Ivy (Hedera helix L.) Flower Nectar and Nectary Ecophysiology. International Journal of Plant Sciences, 167(3), 519–527. https://doi.org/10.1086/501140

Vollmer, M., & Möllmann, K.-P. (2017). Infrared thermal imaging: Fundamentals, research and applications.John Wiley & Sons.

von Arx, M. (2013).Floral humidity and other indicators of energy rewards in pollination biology. Communicative & Integrative Biology, 6(1), e22750. https://doi.org/10.4161/cib.22750

von Arx, M., Goyret, J., Davidowitz, G., & Raguso, R. A. (2012). Floral humidity as a reliable sensory cue for profitability assessment by nectar-foraging hawkmoths. Proceedings of the National Academy of Sciences of the USA, 109, 9471–9476. https://doi.org/10.1073/pnas.1121624109

Wolfin, M. S., Raguso, R. A., Davidowitz, G., & Goyret, J. (2018). Context dependency of in-flight responses by Manduca sexta moths to ambient differences in relative humidity. Journal of Experimental Biology, 221(11). https://doi.org/10.1242/jeb.177774

