## Supplementary material for "The role of petal transpiration in floral humidity generation"

**SUPPLEMENTARY INFORMATION**

**Supplementary information 1: Extended detail of robot sampling procedure**

The method used for measurement of humidity within sample headspace is a modified version of that used by Harrap et al. (2020, 2021). The main text discusses this method and where the procedures differ from that used previously. In this section we recount the protocol used in the current study in further detail.

This method used a Staubli RX 160 robot arm (Pfäffikon, Switzerland) mounted to the floor of a 3.72 m × 3.67 m sampling zone separated by a polycarbonate wall from the rest of the lab, its controller unit and paired PC (the full lab dimensions being 6.11 m × 3.67 m). A humidity probe (DHT-22 humidity probe, Aosong Electronics, Huangpu, China, the 'focal probe') and its Arduino UNO (Adafruit Industries, New York, NY, United States) microcontroller was mounted onto the robot, with the probe attached to a 30cm long bar (see figure 2). Thus, the focal probe is held at a distance from larger moving robot parts. A table (75 cm × 90 cm × 74 cm, width × length × height) was placed within the sampling zone (the area that can be reached by the robot-manipulated probe), and a horticultural tube rack was attached to it, into which prepared samples could be placed ready for sampling. An additional 17cm tall mount was placed on the table, containing a second humidity probe (referred to as the 'background probe') and microcontroller. The background probe was fixed 44.5-54.5cm from any hole in the rack, which we deemed as sufficiently far to not be exposed to humidity of samples. See Harrap et al. (2020) for further details of the robot sampling room, probe mountings and a scale floorplan.

Once a pair of samples were prepared and treatments applied as appropriate (as described in the main text), they were placed in the horticultural tube rack, 15.5cm apart, ready for sampling. Preparation of samples and treatment applications was carried out quickly, and humidity sampling with the robot commenced within one hour of samples being collected. The measurement points along the transects were calculated by the robot as offsets in three-dimensional space (axis directions corresponding to the robot's orientation, see floorplan diagrams in Harrap et al. 2020) relative to a 'transect central point' manually input by operators. Once a pair of samples were ready for humidity sampling and placed in the horticultural tube rack, the focal probe was piloted under manual control to the transect central point of each sample, and this position was input into the robot's memory. This allowed the robot to return to this position (and positions relative to it) autonomously during sampling. Following input of transect central points, and the point 2.5cm away from the background probe for probe-calibration, the robot was then piloted to the 'safe position' a point 50cm above the table clear of samples, and its automated sampling sequence started. The positioning of transect central points for samples is detailed in the main text and figure 3.

All humidity measurements were conducted autonomously by the robot in a set sequence. The robot would randomly select the order in which each sample in a pair would be measured. The focal probe was then moved by the robot arm through the headspace of a sample in a set transect sequence, conducting first an *x* axis transect (horizontal) and then a *z* axis transect (vertical). During these two transects, the arm stopped to take humidity measurements of the sample's headspace at set measurement points along each transect. The exact positions of measurement points relative to the transect central point are detailed in figure 3. Simultaneously to focal probe humidity measurements in the flower headspace, the ambient humidity of the room was measured by the background probe. Following completion of both transects on a sample, the arm then conducted a 'probe-control' step, where the focal probe is moved to a position 2.5 cm away from the background probe, and then both probes simultaneously measure humidity, meaning that both probes are effectively measuring the humidity of the same point in space. This allows us to assess how much background and focal probes differ in their humidity measurements (which can be up to ±5% according to the manufacturer's specifications) and adjust focal probe measurements to account for this allowing calculation of 'corrected focal humidity measurements' (see below, and Harrap et al. 2020). Following a probe control step, the robot then conducted the same transects followed by a probe control step on the other sample in the pair. The robot then repeated this sequence a second time (*x* and *z* transects, then the probe control step) on each sample, maintaining the same sampling order within the pair. Following completion of the last probe control step on the second replicate of the second sample, the arm would return to the 'safe' position and power down. All movements between sample headspaces and the background probe for probe-calibration steps (but not within transect movements) were made *via* the robot's neutral 'safe' position, a point 50cm above the table to prevent sample headspaces being disrupted by motion through its headspace. Throughout the sequence the arm moved at a slow speed, at 3% of the robot's nominal speed (below 200 mm $s^{-1}$).

The transects themselves consisted of an *x* axis transect, across the span of the sample and the *z* axis transect, which moved vertically upwards away from it. In the robot's coordination *x* is the forward-backward horizontal plane, with positive values showing increased distance away from the arm. The *z* axis is the up-down vertical plane, with positive values showing increased vertical height. The robot started each set of transects as follows. The focal probe was moved from the safe position to a point -30mm offset from the sample's transect central point in the robot's coordination. This was the first measurement point in the *x* axis. The arm then moved the focal probe along the *x* axis, through the transect central point to a point +30mm offset from the sample's transect central point. Along this *x* axis transect the robot stopped at measurement points in 10mm intervals. Upon completing the last measurement point on the x axis transect (the point +30mm offset in the *x* axis) the arm was then moved directly to a point +10mm offset in the *z* axis relative to the transect central point. The arm then continued to move upwards along the *z* axis to a point +30mm relative to the transect central point, stopping to also measure humidity at the halfway point between these (+20mm offset in the *z* axis). A full schematic of the measurement points of the transects is given in figure 3.

As detailed in the main text, at each measurement point would stop for 230 seconds during which both probes would take approximately 100 humidity measurements. Measurements by the focal probe would be the 'uncorrected focal relative humidity' ($f_{uncorrected}$), while measurements by the background would be the 'background relative humidity' ($f_{background}$). The robot arm would similarly stop for 230 seconds and take c.100 measurements during the probe control step, where focal and background probes are assumed to measure an area of the same humidity. The measurements collected during the probe control step would be used to correct for slight variation between the two humidity probes in their estimations of the same levels of humidity (up to ±5% according to the manufacturer's specifications), to calculate a 'corrected focal relative humidity' ($f_{corrected}$) reducing this source of inaccuracy in our measurements. Assuming that both probes are measuring a point of the same humidity during the probe-control measurements we compute linear regression parameters, using the MATLAB (2018) 'regression' function, to predict one probe's measurements from the other during the probe control steps. This was later used to obtain adjusted focal humidity measurements using:

$$f_{corrected} = W \cdot f_{uncorrected} + M \quad \text{(S1)}$$

where $W$ and $M$ are, respectively, the slope and intercept parameters obtained from regressing the focal probe measurements against the background probes measurements for the time period of the probe-control measurements. This focal probe correction (Eq. S1) was calculated and applied across each replicate sample of the flowers sampled on the same day (i.e., one set of x and z axis transects on each sample pair). This was done to account for any possibility that the difference between probes may change over time.

From $f_{background}$ and $f_{corrected}$ values $\Delta RH$ values can be calculated (Eq. 1). Mean $\Delta RH$ values for each measurement period (that is the mean of the c.100 measurements taken at each sampling point in the headspace of each replicate on each sample) were then calculated and these were analysed as described in the main text (and described in further detail in supplementary information 2 below).

**Supplementary Information 2: Further detail of models and model simplification procedures**

The full model fitted to the data of the *x* axis humidity transects of each sample type (Dry tube control, Full tube control, *Hedera helix* leaves, *Calystegia silvatica* flowers or *Eschscholzia californica* flowers) is as follows:

$$\begin{aligned}\Delta RH_{xnt} = I_{Cx} + F(i_{Cx}) + [A_{Cx} + F(a_{Cx})]X + [B_{Cx} + F(b_{Cx})]X^2 \\ +U\{I_{Ux} + F(i_{Ux}) + [A_{Ux} + F(a_{Ux})]X + [B_{Ux} + F(b_{Ux})]X^2\} \\ +P\{I_{Px} + F(i_{Px}) + [A_{Px} + F(a_{Px})]X + [B_{Px} + F(b_{Px})]X^2\} + v_{nx} \ .\end{aligned} \tag{S2}$$

Where $\Delta RH_{xnt}$ is the mean *ΔRH* for the measurement period at sampling point $X$, on transect repeat $t$, on flower $n$. $X$ is the *x* axis offset (in mm) of the sampling point (ranging from -30 to +30, in 10mm intervals, see figure 3) relative to the transect central point (which has an x axis offset, $X$, of zero). The replicate transect number is donated by $t$, which can have values of 1 or 2, for the first and second replicate transects of a sample. Parameter $I_{Cx}$ describes the model intercept of samples in the Untreated group in the initial transect. In a similar vein, $A_{Cx}$ and $B_{Cx}$ are parameters that describe the positioning and slope of the *x* axis humidity profile of samples in the Untreated group in the initial transect. Parameters $P$ and $U$ are Boolean operators that allow the model to alter $\Delta RH_{xnt}$ depending on the treatment group samples are in. $U$ denotes whether the sample is in the Unhandled group, where:

$$U = \begin{cases} 0, & sample\ is\ not\ in\ Unhandled\ group \\ 1, & sample\ is\ in\ Unhandled\ group \end{cases}, \tag{S3}$$

and $P$ whether the sample is in the Gel treatment group, where:

$$P = \begin{cases} 0, & sample\ is\ not\ in\ Gel\ group \\ 1, & sample\ is\ in\ Gel\ group \end{cases}. \tag{S4}$$

The action of these Boolean parameters allows several parameters describing differences in humidity profiles relative to the Untreated group to be applied. $I_{Ux}$ and $I_{Px}$ are the change in model intercept, relative to $I_{Cx}$, of samples in the Unhandled and Gel treatment group respectively. Similarly, $A_{Ux}$ and $A_{Px}$ describe the change in positioning of the humidity profiles, relative to $A_{Cx}$, for samples in the Unhandled and Gel treatment group respectively. Lastly, $B_{Ux}$ and $B_{Px}$ describe the change in slopes of the humidity profiles, relative to $B_{Cx}$, for samples in the Unhandled and Gel treatment group respectively. In this way, *via* action of

Boolean parameters $U$ and $P$ and associated parameters, the full model shown in equation S2 allows flowers in different treatment groups to differ in humidity intensity and structure. Flowers may show changes in humidity intensity and structure between replicate transects (Harrap et al., 2020), similarly these changes with replicate transects may differ between treatment groups. Consequentially, for samples in test group "$G$" ($G$ being either $C$, $U$ or $P$ for the Untreated, Unhandled and Gel treatment groups respectively), parameters $i_{Gx}$, $a_{Gx}$ and $b_{Gx}$ modify the effects of $I_{Gx}$, $A_{Gx}$ and $B_{Gx}$ in the second transect, controlled by Boolean operator $F$, where:

$$F = \begin{cases} 0, t = 1 \\ 1, t = 2 \end{cases}. \tag{S5}$$

Thus, parameters $i_{Gx}$, $a_{Gx}$ and $b_{Gx}$ represent the change in humidity intensity, offset and slope (relative to $I_{Gx}$, $A_{Gx}$ and $B_{Gx}$ respectively) in the second transect of samples in test group $G$. Model intercepts are also modified by random factor $v_{nx}$, which represents the change in model intercept, and consequently the intensity of humidity generated, by individual sample $n$ on the *x* axis transect. Further random factors accounting for individual variation in the shape of humidity structure were not included, to simplify models. Here all iterations of to $I_{Gx}$, $A_{Gx}$, $B_{Gx}$, $i_{Gx}$, $a_{Gx}$ and $b_{Gx}$ ($G$ as described above) as well as $v_{nx}$ are parameters to be estimated.

The full model fitted to the data of the *x* axis humidity transects of each sample type is as follows:

$$\begin{aligned} \Delta RH_{znt} = {} & I_{Cz} + F(i_{Cz}) + [B_{Cz} + F(b_{Cz})]\ln(Z) \\ & + U\{I_{Uz} + F(i_{Uz}) + [B_{Uz} + F(b_{Uz})]\ln(Z)\} \\ & + P\{I_{Pz} + F(i_{Pz}) + [B_{Pz} + F(b_{Pz})]\ln(Z)\} + v_{nz}\ . \end{aligned} \tag{S6}$$

Here, parameter $Z$ refers to the sampling point's *z* axis offset relative to the transect central point (figure 3). All other parameters function in the same manner described for respective parameters in the *x* axis model. Here all iterations of to $I_{Gz}$, $B_{Gz}$, $i_{Gz}$ and $b_{Gz}$ ($G$ as described above) as well as $v_{nz}$ are parameters to be estimated.

The first step of analysis was to find the best fitting structure of humidity for each sample type, using models of humidity structure that allowed humidity production to vary as required with treatments, see main text. The full models for the *x* and *z* transects described in Eqs (S2) and (S6), as well as simpler versions of these models, were fitted to the *x* and *z* transect data of each sample type. Simpler models were achieved by removing certain parameters from the full models, done by forcing these parameters to have values of zero. At this stage, eleven *x* axis models and five *z* axis models were compared for each sample. These models are summarized in Tables S1, S2 for the *x* and *z* axis models, respectively. These models were then compared using AIC to select the best models following the guidelines within Richards (2008).

**Table S1:** The models fitted to each sample's x axis transect humidity data, while identifying best humidity structure models. Model name, a model description and the parameters omitted from the full model to create it are given. Additionally, the 'Fullest potential extent of treatment effect' is given, that is the assumed effects of treatment during assessment of humidity structure, these are tested for in the subsequent steps. Note that 'slope' in linear models is influenced by parameters $A_{Gx}$ and $a_{Gx}$ (which influence offset in quadratic models). Use of $G$ in subscript indicates all parameters of that kind (*i.e.* it includes corresponding $C$, $U$ and $P$ subscripted parameters).

| **Model** | **Omitted parameters** | **Model description** | **Fullest potential extent of treatment effect:** (treatment can influence… |
|---|---|---|---|
| m0 | $A_{Gx}, B_{Gx}, a_{Gx}, b_{Gx}, i_{Gx}$ | Flat linear model with no replicate effects | … intercept) |
| m1 | $B_{Gx}, a_{Gx}, b_{Gx}, i_{Gx}$ | Linear model with no replicate effects | … intercept and slope) |
| m2 | $A_{Gx}, a_{Gx}, b_{Gx}, i_{Gx}$ | Quadratic model with no replicate effects | ... intercept and slope) |
| m3 | $a_{Gx}, b_{Gx}, i_{Gx}$ | Quadratic model with x axis offset with no replicate effects | ... intercept, slope and offset) |
| m4 | $A_{Gx}, B_{Gx}, a_{Gx}, b_{Gx}$ | Flat linear model with differing intercepts with replicate transects | ... intercept and differences in intercept with replicate transect) |
| m5 | $B_{Gx}, a_{Gx}, b_{Gx}$ | Linear model with differing intercepts with replicate transects | ... intercept, slope and differences in intercept with replicate transect) |
| m6 | $A_{Gx}, a_{Gx}, b_{Gx}$ | Quadratic model with differing intercepts with replicate transects | ... intercept, slope and differences in intercept with replicate transect) |
| m7 | $a_{Gx}, b_{Gx}$ | Quadratic model with x axis offset with differing intercepts with replicate transects | ... intercept, slope, offset and differences in intercept with replicate transect) |
| m8 | $B_{Gx}, b_{Gx}$ | Linear model with interacting replicate effects | ... intercept and slope and how these change with replicate transect) |
| m9 | $A_{Gx}, a_{Gx}$ | Quadratic model with interacting replicate effects | ... intercept, slope and how these change with replicate transect) |
| m10 | None | The full model (eq. S2). Quadratic model with x axis offset with interacting replicate effects | ... intercept, slope, offset and how these change with replicate transect) |

**Table S2:** The models fitted to each sample's z axis transect humidity data, while identifying best humidity structure models. Model name, a model description and the parameters omitted from the full model to create it are given. Additionally, the 'Fullest potential extent of treatment effect' is given, that is the assumed effects of treatment during assessment of humidity structure, these are tested for in the subsequent steps. Use of $G$ in subscript indicates all parameters of that kind (*i.e.* it includes corresponding $C$, $U$ and $P$ subscripted parameters).

| Model | Omitted parameters | Model description | Fullest potential extent of treatment effect: (treatment can influence… |
|---|---|---|---|
| z0 | $B_{Gz}$, $b_{Gz}$, $i_{Gz}$ | Flat linear model with no replicate effects | … intercept) |
| z1 | $b_{Gz}$, $i_{Gz}$ | Logarithmic model with no replicate effects | … intercept and slope) |
| z2 | $B_{Gz}$, $b_{Gz}$ | Flat linear model with differing intercepts with replicate transects | ... intercept and differences in intercept with replicate transect) |
| z3 | $b_{Gz}$ | Logarithmic model with differing intercepts with replicate transects | … intercept, slope and differences in intercept with replicate transect) |
| z4 | None | The full model (eq. S6). Logarithmic model with interacting replicate effects | … intercept, slope and how these change with replicate transect) |

The second step of the analysis was then to evaluate the effects of treatment on humidity production in each sample type. Here models with different treatment effects were created and fit to data of each sample, by removing these treatment effects parameters form the best fitting model of humidity structure identified in the previous step. Five different kinds of treatment effects models were created as summarized in Table S3. Where the best fitting model of humidity structure for a sample, as selected above, did not include changes in humidity with replicate effects the T4 and Tz4 models were not fitted to the data (as there are no replicate effects for treatment to alter). Depending on the best performing humidity structure model, either the T3 and Tz3 or the T4 and Tz4 model represent the 'full' model selected in the humidity structure step (it was possible for T1 and Tz1 to be the 'full' model but this was not the case in any of our samples, see main text).

**Table S3**: the models fitted to each sample's humidity data during assessment of treatment effects. Models fitted to both *x* axis and *z* axis data are given. Given are the model names, a description and the parameters omitted. Note, that parameters omitted are those omitted when they are present in the best fitting model of humidity structure identified previously, as discussed in the above and main text. In addition to these in samples that underwent three treatments further variant of these models were created to further explore differences between treatment groups (detailed below).

| | **Model** | **Omitted parameters** (when they are present in best fitting structure model) | **Model description** |
|---|---|---|---|
| x axis models | *T0* | $I_{Ux}$, $I_{Px}$, $A_{Ux}$, $A_{Px}$, $B_{Ux}$, $B_{Px}$, $i_{Ux}$, $i_{Px}$, $a_{Ux}$, $a_{Px}$, $b_{Ux}$, $b_{Px}$ | no treatment effect on *x* axis humidity production |
| | *T1* | $A_{Ux}$, $A_{Px}$, $B_{Ux}$, $B_{Px}$, $i_{Ux}$, $i_{Px}$, $a_{Ux}$, $a_{Px}$, $b_{Ux}$, $b_{Px}$ | treatment effect on *x* axis humidity intensity only |
| | *T2* | $I_{Ux}$, $I_{Px}$, $i_{Ux}$, $i_{Px}$, $a_{Ux}$, $a_{Px}$, $b_{Ux}$, $b_{Px}$ | treatment effect on *x* axis humidity structure only |
| | *T3* | $i_{Ux}$, $i_{Px}$, $a_{Ux}$, $a_{Px}$, $b_{Ux}$, $b_{Px}$ | treatment effect on both *x* axis humidity intensity and structure |
| | *T4* | None | treatment effect on both *x* axis humidity intensity and structure, as well as how humidity changes with replicate effects |
| z axis models | *Tz0* | $I_{Uz}$, $I_{Pz}$, $B_{Uz}$, $B_{Pz}$, $i_{Uz}$, $i_{Pz}$, $b_{Uz}$, $b_{Pz}$ | no effect treatment on *z* axis humidity production |
| | *Tz1* | $B_{Uz}$, $B_{Pz}$, $i_{Uz}$, $i_{Pz}$, $b_{Uz}$, $b_{Pz}$ | treatment effect on *z* axis humidity intensity only |
| | *Tz2* | $I_{Uz}$, $I_{Pz}$, $i_{Uz}$, $i_{Pz}$, $b_{Uz}$, $b_{Pz}$ | treatment effect on *z* axis humidity structure only |
| | *Tz3* | $i_{Uz}$, $i_{Pz}$, $b_{Uz}$, $b_{Pz}$ | treatment effect on both *z* axis humidity intensity and structure |
| | *Tz4* | None | treatment effect on both *z* axis humidity intensity and structure, as well as how humidity changes with replicate effects |

In samples that underwent three treatment groups a further set of these treatment effect models were fitted to the data to further assess differences between treatment groups, see main text. Models lacking further subscript in their identifiers, were as described in table S3, allowing the three groups to differ in all treatment effects present in the model. Models with the 'TCwP' subscript where Untreated and Gel treatments are grouped together were created by forcing parameter $P$ to always equal 0 (effectively removing parameters $I_{Px}$, $A_{Px}$, $B_{Px}$, $i_{Px}$, $a_{Px}$ and $b_{Px}$, when present in the *x* axis model, and $I_{Pz}$, $B_{Pz}$, $i_{Pz}$ and $b_{Pz}$, when present in the *z* axis model). This meant humidity of samples in the Untreated and Gel group would be determined by the same parameters. The 'TCwU' models, where Untreated and Unhandled treatments are grouped together would be similarly created by forcing parameter $U$ to always equal 0, similarly removing the parameters associated with the Unhandled group and determining humidity of both groups by common parameters. The 'TPwU' models, where Gel and Unhandled treatments are grouped together were created by forcing parameter $P$ to always equal 0, but substituting $U$ with $U^{alt}$, where:

$$U^{alt} = \begin{cases} 0, & sample\ is\ in\ Untreated\ group \\ 1, & sample\ is\ in\ Unhandled\ or\ Gel\ group \end{cases}. \quad \text{(S7)}$$

This removes the parameters associated with $P$ but now allows Gel group humidity be determined by the same parameters as the Unhandled group. Note that, the *T0* and *Tz0* models (table S3) describe no treatment effects, thus there were no further variants of these models. These treatment effect models were then compared using AIC to select the best models following the guidelines within Richards (2008).

Datafiles and code for the analyses described in this study and figure generation is available within Harrap and Rands (2021).

### Supplementary information 3: Summary value calculation

Humidity intensity summary values $X_t^{max}$ (the position of the mean peak in humidity production over the x axis transect, value of $X$ relative to the transect central point) and $\Delta RH_x^{max}$ (the average peak in humidity production over the *x* axis transect) were calculated according to the best fitting treatment model for each treatment of each sample and each replicate transect when it was included in the best fitting model. These calculations were the same as those used to produce these values in Harrap et al. (2020, 2021) but with adjustments for the presence of different treatment groups.

Where the best fitting *x* axis treatment model was flat in structure, $X_t^{max}$ for each treatment of each sample and each replicate transect was taken as 0, although any value of $X$ would have the same $\Delta RH$ value and thus be a suitable point for the 'peak' humidity. Where the best fitting *x* axis treatment model was of a linear structure $X_t^{max}$ was

$$X_t^{max} \begin{cases} -30, A_{Cx} + F(a_{Cx}) + U(A_{Ux} + F(a_{Ux})) + P(A_{Px} + F(a_{Px})) < 0 \\ +30, A_{Cx} + F(a_{Cx}) + U(A_{Ux} + F(a_{Ux})) + P(A_{Px} + F(a_{Px})) > 0 \end{cases} . \quad \text{(S8)}$$

For quadratic models, the $X_t^{max}$ for each treatment of each sample and each replicate transect could be calculated as follows:

$$X_t^{max} = -\frac{A_{Cx} + F(a_{Cx}) + U(A_{Ux} + F(a_{Ux})) + P(A_{Px} + F(a_{Px}))}{2(B_{Cx} + F(b_{Cx}) + U(B_{Ux} + F(b_{Ux})) + P(B_{Px} + F(b_{Px})))} . \quad \text{(S9)}$$

In both equations S8 and S9: values of $F$ are 0 or 1 depending on the replicate transect for which values are being calculated (see equation S5); values for $U$ and $P$ are 0 and 1 depending on the treatment groups for which values are being calculated (see equations S3, S4 and S7); and all other parameters are the values estimated by the best fitting treatment effect model of the sample. Note that if the best fitting treatment effect model were a 'TPwU' model $U^{alt}$ would be used in place of $U$ in equations S8 and S9.

With $X_t^{max}$ calculated $\Delta RH_x^{max}$ for each treatment of each sample and each replicate transect using:

$$\begin{aligned} \Delta RH_x^{max} = {} & I_{Cx} + F(i_{Cx}) + [A_{Cx} + F(a_{Cx})]X_t^{max} + [B_{Cx} + F(b_{Cx})](X_t^{max})^2 \\ & +U\{I_{Ux} + F(i_{Ux}) + [A_{Ux} + F(a_{Ux})]X_t^{max} + [B_{Ux} + F(b_{Ux})](X_t^{max})^2\} \\ & +P\{I_{Px} + F(i_{Px}) + [A_{Px} + F(a_{Px})]X_t^{max} + [B_{Px} + F(b_{Px})](X_t^{max})^2\} . \end{aligned}$$

(S10)

Where: $X_t^{max}$ is the $X_t^{max}$ value appropriate to the treatment group, replicate transect and sample for which values are being calculated; values of $F$ are 0 or 1 depending on the replicate transect for which values are being calculated (see equation S5); values for $U$ and $P$ are 0 and 1 depending on the treatment groups for which values are being calculated (see equations S3, S4 and S7); and all other parameters are the values estimated by the best fitting treatment effect model of the sample. Note that if the best fitting treatment effect model were a 'TPwU' model $U^{alt}$ would be used in place of $U$ in equations S8 and S9.

Code for calculation of summary values can be found within the datafiles associated with this research (Harrap and Rands, 2021).

**Supplementary Information 4: AIC tables**

Presented below are the AIC tables relating to the analysis of humidity structure and treatment effects of each tube control, *Hedera helix* leaves and flowers of *Calystegia silvatica* and *Eschscholzia californica*. Provided for each transect of each species and control are the AIC tables for selecting the best model for humidity structure assuming treatment effects followed by AIC tables for model comparisons assessing effects of treatments.

Given for each model is its name 'Model', degrees of freedom 'df', AIC values 'AIC', and the difference between the model's AIC value and that of the model with the lowest AIC in each set. Bold models indicate the best or (where appropriate) comparable best models as per the guidelines laid out by Richards (2008). For details of analysis see Supplementary Information 2.

In humidity structure tables humidity structure of each model is codified in column 'struc': '*F*' indicating a flat structure, '*L*' a linear (*x* axis) or log-linear (*z* axis) structure, and '*Q*' an quadratic structure (*x* axis only) and '*oQ*' an offset quadratic structure (*x* axis only). The presence of subset letters in column '*s*' indicates the model allows changes in humidity with replicate transects: a subscript '*r*' indicating replicate transect effects on the model intercept, and a subscript '*e*' indicating replicate transects influence both intercept and structure of the model (interacting effects).

In the treatment effects tables, the effects of treatment is codified in column 'treat': '*n*' no effect of treatment, '*i*' treatment effects model intercepts, '*s*' treatment effects model structure, '*is*' treatment effects both model intercept and structure, '*r*' treatment effects model intercept, structure and replicate transect effects. Note if the best humidity structure model does not include replicate effects treatment's effect on replicate effects is not tested, thus that model is absent.

Where more than two treatments are present, each models grouping of treatment effects is indicated in the subscript of the treatment effects model name exactly as described in the main text: a lack of a subscript entry indicating all treatments differ; 'TCwP', Untreated and Gel treatments are grouped together; 'TCwU', Untreated and Unhandled treatments are grouped together; 'TPwU', Gel treatment and Unhandled treatments are grouped together.

### A) Dry tube control

Dry tube control - *x* axis models

Humidity structure

| Model | struc | df | AIC | ΔAIC |
|---|---|---|---|---|
| m0 | *F* | 4 | -214.598 | 73.061 |
| m1 | *L* | 6 | -260.357 | 27.302 |
| m2 | *Q* | 6 | -236.371 | 51.288 |
| **m3** | ***oQ*** | **8** | **-287.659** | **0.000** |
| m4 | $F_r$ | 6 | -211.436 | 76.223 |
| m5 | $L_r$ | 8 | -257.365 | 30.295 |
| m6 | $Q_r$ | 8 | -233.293 | 54.366 |
| m7 | $oQ_r$ | 10 | -284.791 | 2.868 |
| m8 | $L_e$ | 10 | -254.969 | 32.690 |
| m9 | $Q_e$ | 10 | -229.307 | 58.352 |
| m10 | $oQ_e$ | 14 | -278.611 | 9.048 |

Treatment effects

| Model | treat | df | AIC | ΔAIC |
|---|---|---|---|---|
| **T0** | ***n*** | **5** | **-292.108** | **0.000** |
| T1 | *i* | 6 | -290.786 | 1.322 |
| T2 | *s* | 7 | -288.660 | 3.449 |
| T3 | *is* | 8 | -287.659 | 4.449 |

Dry tube control - *z* axis models

Humidity structure

| Model | struc | df | AIC | ΔAIC |
|---|---|---|---|---|
| z0 | *F* | 4 | -216.469 | 67.352 |
| **z1** | ***L*** | **6** | **-283.821** | **0.000** |
| z2 | $F_r$ | 6 | -214.256 | 69.564 |
| z3 | $L_r$ | 8 | -282.708 | 1.113 |
| z4 | $L_e$ | 10 | -281.203 | 2.618 |

Treatment effects

| Model | treat | df | AIC | ΔAIC |
|---|---|---|---|---|
| **Tz0** | ***n*** | **4** | **-287.441** | **0.000** |
| Tz1 | *i* | 5 | -285.741 | 1.700 |
| Tz2 | *s* | 5 | -285.656 | 1.785 |
| Tz3 | *is* | 6 | -283.821 | 3.620 |

### B) Full tube control

Full tube control - *x* axis models

Humidity structure

| Model | struct | df | AIC | ΔAIC |
|---|---|---|---|---|
| m0 | *F* | 4 | -85.221 | 160.883 |
| m1 | *L* | 6 | -81.653 | 164.451 |
| m2 | *Q* | 6 | -236.365 | 9.738 |
| m3 | *oQ* | 8 | -233.029 | 13.074 |
| m4 | $F_r$ | 6 | -90.100 | 156.003 |
| m5 | $L_r$ | 8 | -86.543 | 159.561 |
| **m6** | $\boldsymbol{Q_r}$ | **8** | **-246.103** | **0.000** |
| m7 | $oQ_r$ | 10 | -242.793 | 3.310 |
| m8 | $L_e$ | 10 | -83.325 | 162.778 |
| m9 | $Q_e$ | 10 | -242.132 | 3.971 |
| m10 | $oQ_e$ | 14 | -236.043 | 10.060 |

Treatment effects

| Model | treat | df | AIC | ΔAIC |
|---|---|---|---|---|
| T0 | *n* | 5 | -239.922 | 6.928 |
| **T1** | ***i*** | **6** | **-246.851** | **0.000** |
| **T2** | ***s*** | **6** | **-243.889** | **2.961** |
| T3 | *is* | 7 | -244.911 | 1.940 |
| T4 | *r* | 8 | -246.103 | 0.747 |

Full tube control - *z* axis models

Humidity structure

| Model | struc | df | AIC | ΔAIC |
|---|---|---|---|---|
| z0 | *F* | 4 | -389.577 | 39.238 |
| z1 | *L* | 6 | -415.533 | 13.282 |
| z2 | $F_r$ | 6 | -400.467 | 28.348 |
| **z3** | $\boldsymbol{L_r}$ | **8** | **-428.814** | **0.000** |
| z4 | $L_e$ | 10 | -425.591 | 3.223 |

Treatment effects

| Model | treat | df | AIC | ΔAIC |
|---|---|---|---|---|
| **Tz0** | ***n*** | **5** | **-426.051** | **2.764** |
| Tz1 | *i* | 6 | -425.679 | 3.136 |
| Tz2 | *s* | 6 | -425.154 | 3.660 |
| Tz3 | *is* | 7 | -425.530 | 3.285 |
| Tz4 | *r* | 8 | -428.814 | 0.000 |

#### C) *Hedera helix* leaves

*Hedera helix* leaves - *x* axis models

Humidity structure

| Model | struc | df | AIC | ΔAIC |
|---|---|---|---|---|
| m0 | *F* | 4 | -52.854 | 61.707 |
| **m1** | ***L*** | **6** | **-114.560** | **0.000** |
| m2 | *Q* | 6 | -51.306 | 63.254 |
| m3 | *oQ* | 8 | -114.450 | 0.110 |
| m4 | $F_r$ | 6 | -50.252 | 64.308 |
| m5 | $L_r$ | 8 | -112.773 | 1.787 |
| m6 | $Q_r$ | 8 | -48.728 | 65.832 |
| m7 | $oQ_r$ | 10 | -112.724 | 1.836 |
| m8 | $L_e$ | 10 | -109.924 | 4.636 |
| m9 | $Q_e$ | 10 | -45.010 | 69.551 |
| m10 | $oQ_e$ | 14 | -106.362 | 8.198 |

Treatment effects

| Model | treat | df | AIC | ΔAIC |
|---|---|---|---|---|
| T0 | *n* | 4 | -72.300 | 42.260 |
| T1 | *i* | 5 | -83.378 | 31.183 |
| T2 | *s* | 5 | -103.483 | 11.078 |
| **T3** | ***is*** | **6** | **-114.560** | **0.000** |

*Hedera helix* leaves - *z* axis models

Humidity structure

| Model | struc | df | AIC | ΔAIC |
|---|---|---|---|---|
| z0 | *F* | 4 | -180.504 | 31.128 |
| **z1** | ***L*** | **6** | **-211.632** | **0.000** |
| z2 | $F_r$ | 6 | -177.346 | 34.286 |
| z3 | $L_r$ | 8 | -208.941 | 2.690 |
| z4 | $L_e$ | 10 | -205.482 | 6.149 |

Treatment effects

| Model | treat | df | AIC | ΔAIC |
|---|---|---|---|---|
| Tz0 | *n* | 4 | -172.977 | 38.655 |
| Tz1 | *i* | 5 | -183.064 | 28.568 |
| Tz2 | *s* | 5 | -172.265 | 39.366 |
| **Tz3** | ***is*** | **6** | **-211.632** | **0.000** |

### D) *Calystegia silvatica* flowers

*Calystegia silvatica* - *x* axis models

Humidity structure

| Model | struc | df | AIC | ΔAIC |
|---|---|---|---|---|
| m0 | *F* | 5 | 943.182 | 515.088 |
| m1 | *L* | 8 | 880.666 | 452.572 |
| m2 | *Q* | 8 | 573.407 | 145.313 |
| m3 | *oQ* | 11 | 448.898 | 20.804 |
| m4 | $F_r$ | 8 | 937.323 | 509.229 |
| m5 | $L_r$ | 11 | 873.430 | 445.336 |
| m6 | $Q_r$ | 11 | 557.706 | 129.612 |
| **m7** | $\boldsymbol{oQ_r}$ | **14** | **428.094** | **0.000** |
| m8 | $L_e$ | 14 | 878.297 | 450.203 |
| m9 | $Q_e$ | 14 | 559.588 | 131.494 |
| m10 | $oQ_e$ | 20 | 432.643 | 4.549 |

Treatment effects

| Model | treat | df | AIC | ΔAIC |
|---|---|---|---|---|
| T0 | *n* | 6 | 577.364 | 149.270 |
| T1 | *i* | 8 | 545.375 | 117.281 |
| T2 | *s* | 10 | 495.365 | 67.271 |
| T3 | *is* | 12 | 434.578 | 6.483 |
| **T4** | ***r*** | **14** | **428.094** | **0.000** |
| T1$_{TCwP}$ | *i* | 7 | 577.969 | 149.875 |
| T2$_{TCwP}$ | *s* | 8 | 574.628 | 146.534 |
| T3$_{TCwP}$ | *is* | 9 | 575.783 | 147.689 |
| T4$_{TCwP}$ | *r* | 10 | 577.686 | 149.592 |
| T1$_{TCwU}$ | *i* | 7 | 546.587 | 118.493 |
| T2$_{TCwU}$ | *s* | 8 | 512.369 | 84.275 |
| T3$_{TCwU}$ | *is* | 9 | 457.538 | 29.444 |
| T4$_{TCwU}$ | *r* | 10 | 450.905 | 22.810 |
| T1$_{TPwU}$ | *i* | 7 | 561.116 | 133.022 |
| T2$_{TPwU}$ | *s* | 8 | 523.866 | 95.772 |
| T3$_{TPwU}$ | *is* | 9 | 485.177 | 57.083 |
| T4$_{TPwU}$ | *r* | 10 | 480.456 | 52.362 |

*Calystegia silvatica* - *z* axis models

Humidity structure

| Model | struc | df | AIC | ΔAIC |
|---|---|---|---|---|
| z0 | *F* | 5 | -584.808 | 26.070 |
| z1 | *L* | 8 | -599.768 | 11.111 |
| z2 | $F_r$ | 8 | -594.901 | 15.977 |
| **z3** | $\boldsymbol{L_r}$ | **11** | **-610.878** | **0.000** |
| z4 | $L_e$ | 14 | -606.597 | 4.281 |

Treatment effects

| Model | treat | df | AIC | ΔAIC |
|---|---|---|---|---|
| Tz0 | *n* | 5 | -571.921 | 40.007 |
| Tz1 | *i* | 7 | -586.770 | 25.159 |
| Tz2 | *s* | 7 | -571.030 | 40.899 |
| Tz3 | *is* | 9 | -603.945 | 7.983 |
| Tz4 | *r* | 11 | -610.878 | 1.050 |
| Tz1$_{TCwP}$ | *i* | 6 | -571.293 | 40.636 |
| Tz2$_{TCwP}$ | *s* | 6 | -570.035 | 41.893 |
| Tz3$_{TCwP}$ | *is* | 7 | -571.550 | 40.378 |
| Tz4$_{TCwP}$ | *r* | 8 | -577.714 | 34.215 |
| Tz1$_{TCwU}$ | *i* | 6 | -587.920 | 24.009 |
| Tz2$_{TCwU}$ | *s* | 6 | -572.783 | 39.146 |
| **Tz3$_{TCwU}$** | ***is*** | **7** | **-606.585** | **5.344** |
| Tz4$_{TCwU}$ | *r* | 8 | -611.928 | 0.000 |
| Tz1$_{TPwU}$ | *i* | 6 | -578.784 | 33.145 |
| Tz2$_{TPwU}$ | *s* | 6 | -571.398 | 40.531 |
| Tz3$_{TPwU}$ | *is* | 7 | -586.308 | 25.620 |
| Tz4$_{TPwU}$ | *r* | 8 | -584.311 | 27.617 |

### E) *Eschscholzia californica* flowers

*Eschscholzia californica* - *x* axis models

Humidity structure

| Model | struc | df | AIC | ΔAIC |
|---|---|---|---|---|
| m0 | *F* | 5 | 1391.126 | 379.294 |
| m1 | *L* | 8 | 1362.371 | 350.539 |
| m2 | *Q* | 8 | 1065.809 | 53.977 |
| **m3** | ***oQ*** | **11** | **1011.832** | **0.000** |
| m4 | $F_r$ | 8 | 1394.400 | 382.568 |
| m5 | $L_r$ | 11 | 1365.491 | 353.659 |
| m6 | $Q_r$ | 11 | 1067.190 | 55.358 |
| m7 | $oQ_r$ | 14 | 1012.750 | 0.918 |
| m8 | $L_e$ | 14 | 1371.292 | 359.460 |
| m9 | $Q_e$ | 14 | 1070.546 | 58.714 |
| m10 | $oQ_e$ | 20 | 1021.486 | 9.654 |

Treatment effects

| Model | treat | df | AIC | ΔAIC |
|---|---|---|---|---|
| T0 | *n* | 5 | 1071.433 | 59.601 |
| T1 | *i* | 7 | 1041.921 | 30.089 |
| T2 | *s* | 9 | 1067.541 | 55.709 |
| **T3** | ***is*** | **11** | **1011.832** | **0.000** |
| $T1_{TCwP}$ | *i* | 6 | 1073.381 | 61.549 |
| $T2_{TCwP}$ | *s* | 7 | 1075.175 | 63.343 |
| $T3_{TCwP}$ | *is* | 8 | 1077.132 | 65.300 |
| $T1_{TCwU}$ | *i* | 6 | 1050.285 | 38.453 |
| $T2_{TCwU}$ | *s* | 7 | 1065.285 | 53.453 |
| $T3_{TCwU}$ | *is* | 8 | 1023.945 | 12.113 |
| $T1_{TPwU}$ | *i* | 6 | 1047.518 | 35.686 |
| $T2_{TPwU}$ | *s* | 7 | 1067.956 | 56.124 |
| $T3_{TPwU}$ | *is* | 8 | 1023.300 | 11.468 |

*Eschscholzia californica* – *z* axis models

Humidity structure

| Model | struc | df | AIC | ΔAIC |
|---|---|---|---|---|
| z0 | *F* | 5 | -814.850 | 78.223 |
| **z1** | ***L*** | **8** | **-891.527** | **1.547** |
| z2 | $F_r$ | 8 | -814.796 | 78.278 |
| z3 | $L_r$ | 11 | -893.074 | 0.000 |
| z4 | $L_e$ | 14 | -888.650 | 4.423 |

Treatment effects

| Model | treat | df | AIC | ΔAIC |
|---|---|---|---|---|
| Tz0 | *n* | 4 | -840.596 | 54.477 |
| Tz1 | *i* | 6 | -859.054 | 36.020 |
| Tz2 | *s* | 6 | -840.169 | 54.905 |
| Tz3 | *is* | 8 | -891.527 | 3.546 |
| $Tz1_{TCwP}$ | *i* | 5 | -841.604 | 53.469 |
| $Tz2_{TCwP}$ | *s* | 5 | -838.950 | 56.123 |
| $Tz3_{TCwP}$ | *is* | 6 | -845.171 | 49.903 |
| $Tz1_{TCwU}$ | *i* | 5 | -860.847 | 34.227 |
| $Tz2_{TCwU}$ | *s* | 5 | -842.119 | 52.955 |
| **$Tz3_{TCwU}$** | ***is*** | **6** | **-895.074** | **0.000** |
| $Tz1_{TPwU}$ | *i* | 5 | -846.588 | 48.485 |
| $Tz2_{TPwU}$ | *s* | 5 | -839.696 | 55.378 |
| $Tz3_{TPwU}$ | *is* | 6 | -858.177 | 36.896 |

**References cited in Supplementary Material**

Harrap, M.J.M., de Ibarra, N.H., Knowles, H.D., Whitney, H.M., Rands, S.A., 2021. Bumblebees can detect floral humidity. bioRxiv. https://doi.org/10.1101/2021.03.19.436119

Harrap, M.J.M., Hempel de Ibarra, N., Knowles, H.D., Whitney, H.M., Rands, S.A., 2020. Floral humidity in flowering plants: a preliminary survey. Frontiers in Plant Science 11, 249. https://doi.org/10.3389/fpls.2020.00249

Harrap, M.J.M., Rands, S.A., 2021. Data from "The role of transpiration in floral humidity generation." Figshare Database. https://doi.org/10.6084/m9.figshare.14350547

MATLAB, 2018. version 9.7.0.1190202 (R2018b). The MathWorks Inc., Natick, Massachusetts.

Richards, S.A., 2008. Dealing with overdispersed count data in applied ecology. J Appl Ecol 45, 218–227. https://doi.org/10.1111/j.1365-2664.2007.01377.x
